# Node-specific amplitude alignment and finite response capacity define dynamic phase balancing in a reduced sarcomere-chain model

**DOI:** 10.64898/2026.07.20.739477

**Authors:** Seine A. Shintani

## Abstract

During hyperthermal sarcomeric oscillations (HSOs), serial sarcomeres differ in amplitude and phase. Their fast length changes can cancel in total length, while changing relative amplitudes can move the best phase arrangement. Each model node represented one sarcomere. I tested two requirements: correct amplitude-to-sarcomere pairing and timely phase adjustment. Lower normalized residuals meant better cancellation; subtracting the minimum set by the current amplitudes isolated the part that phase adjustment could remove.

Correct pairing reduced the 95th percentile of this avoidable part relative to amplitude-blind or misassigned inputs in all 20 prespecified conditions. Dynamic adjustment outperformed the best of 13 fixed arrangements in all 12 conditions with target periods of at least four HSO cycles, but only 2/8 faster conditions. A ratio of target-change time to model response time almost perfectly ranked dynamic wins above fixed wins in held-out and speed-limit tests (AUC 0.976 and 0.996), although the best cutoff differed among condition families.

Replaying measured amplitude histories from five sarcomeres in seven cardiomyocytes gave a median avoidable residual of 0.000802, about 1/78 of the lowest fixed-strategy median, and out-performed three fixed strategies in 7/7 cells. Returning each history to its source sarcomere gave the lowest residual among all 120 assignments in every cell. Observed phase motion favored the predicted direction relative to the circular-shift median in 5/7 cells; the prespecified cell-level test gave *P* = 0.1094. Within this reduced model, dynamic phase balancing requires correct node-specific information and sufficient response time. The study establishes this conditional model capacity and identifies native phase dynamics as the next mechanochemical test.

**Significance:** Serial sarcomeres can partly cancel one another’s fast length changes when their oscillations occur at different phases. When local amplitudes differ, the best-cancelling phase pattern depends on which amplitude belongs to which sarcomere, and changes in their relative pattern can move the target. This model separates two requirements: locating the pattern from node-specific amplitudes and following it with a response that has finite speed and delay. Simulations and replay of seven previously recorded cardiomyocytes show when dynamic adjustment outperforms fixed phases and provide concrete tests for future mechanochemical models.

## 1 Introduction

Sarcomeres, the repeating contractile units arranged end to end within a myofibril, operate in mechanically connected arrays, yet adjacent units can shorten and lengthen at different times. When two units move in opposite directions, their length changes partly cancel in the chain total; when they move together, the changes add. The collective question is therefore how local amplitudes and phases combine to determine the fast fluctuation across the serial array. Measurements at single-sarcomere resolution have shown how spatial averaging conceals this local motion[1, 2]. Above approximately 38^◦^C, neonatal-rat cardiomyocytes also exhibit hyperthermal sarcomeric oscillations (HSOs), a fast calcium-independent length oscillation superimposed on slower calcium-dependent beating[3]. Beat-scale synchrony supports coordinated ventricular pumping[4]; the faster HSO component poses a different collective problem because its summed length signal depends on the amplitudes and phases of the participating sarcomeres.

A 2020 study combined five-sarcomere measurements with a 40-half-sarcomere mechanochemical model, which links molecular states, force, and length, and showed that asynchronous local length changes can compensate in total length and model tension[5]. A 2022 analysis then showed that HSO amplitudes display chaotic properties while oscillation cycles remain comparatively constant[6]. The 2026 BPPB analysis of the same seven-cell series found that phase relationships usually changed between one neighboring pair at a time and that the effect of a local phase change on total length depended on the surrounding phases[7]. These studies establish amplitude heterogeneity, phase-offset compensation, and local phase rearrangement as the biological starting point. They leave open a quantitative question: when the relative pattern of local amplitudes changes, what phase arrangement reduces the summed fast signal, and how quickly must the system respond to remain near that arrangement?

Oscillator theory has established that a group can reduce its collective output by distributing phases around the cycle rather than moving in synchrony; an evenly spaced arrangement is often called a splay state[8–10]. Control studies show how feedback can reshape these phase relations when responses are delayed or limited in speed[11–14], while muscle models add serial mechanics and local or global coupling[15–17]. Equal phase spacing guarantees cancellation when local amplitudes are equal, whereas unequal amplitudes generally favor another arrangement. A useful picture is a set of arrows: amplitude sets each arrow’s length and phase sets its direction. The local oscillations cancel when the arrows close into a polygon. When the relative arrow lengths change, the directions that close the polygon can also change, so the best-cancelling phase arrangement can move over time.

I separated two requirements. First, the response must know the amplitude belonging to each node, because those amplitudes specify the phase arrangement that cancels the summed signal most strongly at that moment. Second, the phases must move toward that arrangement before changes in relative amplitudes move it elsewhere. I refer to the first requirement as target definition and the second as target tracking. Lag, relaxation, and a relative-phase speed limit determine the available tracking capacity.

I tested target definition by keeping the amplitudes used to calculate the summed signal unchanged while removing or reassigning only the amplitude information supplied to the phase response. I tested target tracking by comparing the dynamic response with 13 fixed phase arrangements across environmental timescales, delays, and speed limits, including parameter combinations reserved from model development. Finally, I replayed amplitude histories from five sarcomeres in each of seven previously recorded cells and tested all 120 possible within-cell assignments. Throughout the study, the endpoint is the summed HSO-band length signal. Force, tension, mechanical work, and energetic cost remain endpoints for a future mechanochemical model.

## 2 Materials and methods

### 2.1 Study design and experimental provenance

The study combined analytical results, reduced-model simulations, and reanalysis of previously reported recordings. The recording-based analyses used an 8.0-s segment during the HSO state from five serial-sarcomere length traces in each of seven neonatal-rat cardiomyocytes described in the source studies[3, 5]. Sarcomere-length image sequences were acquired at a nominal 500 fps, as reported previously[5]. The cell was the independent biological unit. Time points, sarcomeres, cycles, parameter combinations, simulation seeds, and within-cell assignments served as repeated observations or computational conditions.

The original experiments were approved by the Animal Experiment Committee of the Faculty of Science, The University of Tokyo, and were conducted under the university’s animal-experiment guidelines. Cell preparation, local heating, sarcomere tracking, and animal procedures are reported in the original publications[3, 5]. The present study consists entirely of mathematical and computational analyses of those existing records.

In this article, a causal model update means that the simulated phase response used only the current and past supplied amplitudes. Empirical amplitude and phase histories were extracted offline. Forward– backward filtering removes filter-induced phase delay by processing the signal in both time directions[18], and the Hilbert analytic signal represents a band-limited oscillation by an instantaneous amplitude and phase[19]. The replay therefore quantifies a conditional capacity: the collective output achievable when the declared phase-adjustment rule receives amplitudes extracted from the entire selected 8.0-s analysis segment. A separate one-sided analysis tests how amplitude estimation from present and past samples changes that capacity.

### 2.2 Weighted collective length-signal residual

Over one HSO cycle, the fast length component at model node *i* in a chain of *N* nodes, each representing one sarcomere, was written as

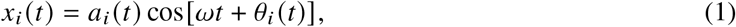

where *a*_*i*_ *(t)* ≥ 0 is the effective amplitude, *0*_*i*_ *(t)* is the relative phase, and *ω* is the common HSO angular frequency. The corresponding weighted phasor—an arrow in the complex plane whose length represents amplitude and whose direction represents phase—and normalized residual were

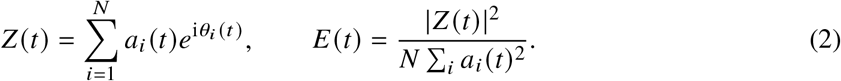

Here *E* measures the part of local HSO-band motion that remains after the node signals are summed and normalized. The normalization keeps *E* between zero and one. A value of zero means complete cancellation, whereas a larger value means that more fast oscillation remains in the collective length signal. The instantaneous minimum over freely selectable phases is the weighted-polygon bound

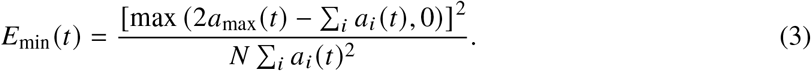

Here *a*_max_ is the largest current node amplitude. The bound is zero whenever the weighted vectors can close a polygon; otherwise, the largest vector leaves an unavoidable residual. The difference *E* − *E*_min_ separates the residual caused by the current phase arrangement, which can in principle be reduced by changing phases, from the residual imposed by the amplitudes themselves. In the equations below, Re and Im denote real and imaginary parts, the superscript denotes complex conjugation, an overdot * denotes a time derivative, and ‖ ·‖_2_ denotes Euclidean length.

The primary endpoint was the 95th percentile of the residual above this instantaneous best value, *Q*_0.95_ [*E* −*E*_min_]. This is the value that the avoidable residual stayed below for 95% of the evaluation window; a smaller value means that the response prevented large episodes of incomplete cancellation. I also compared *Q*_0.95_ (*E)* with fixed phase arrangements. In sensitivity tests, fixed phases were optimized separately for mean *E* or for the mean of the worst 5% of observations, CVaR_0.95_ (*E)*. Difference endpoints were primary because ratios become unstable when the fixed value approaches zero.

For fixed amplitudes, the phase gradient is

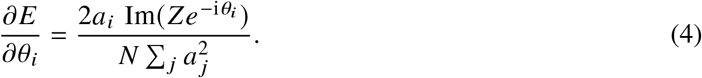

An unconstrained gradient response, 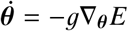, then satisfies

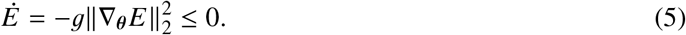

This identity shows that, for fixed amplitudes and unconstrained phase motion, movement opposite to the gradient drives *E* downward. It supplies a best-improvement direction against which the delayed and speed-limited response can be compared.

### 2.3 A moving target and finite phase response

Writing *w*_*i*_ = *a*_*i*_ */*‖***a***‖ _2_ and 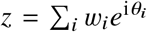 gives *E* = |*z*|^2^ *N*. Its exact derivative separates the effect of changing relative amplitudes from the effect of realized phase motion:

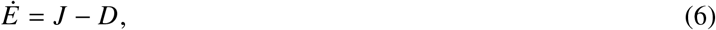

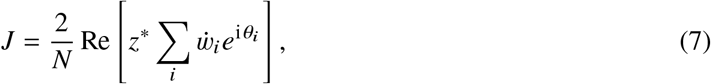

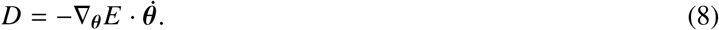

Positive *J* raises the residual as changing relative amplitudes move the best-cancelling configuration, whereas positive *D* records how much the realized phase motion reduces the residual. In plain terms, Eq. (6) describes a race between movement of the target and movement toward the target.

Synthetic amplitudes were generated from a common periodic calcium-like input *C* (*t*), a common load-like input *L* (*t*), and node-specific response parameters:

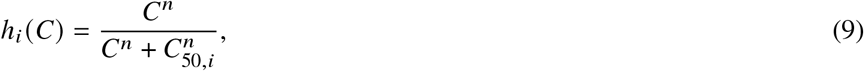

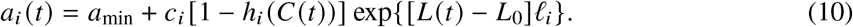

Here *C (t)* and *L (t)* are dimensionless synthetic drivers that create a changing amplitude pattern. The Hill function *h*_*i*_ has exponent *n* and node-specific half-response value *C*_50,*i*_. The remaining terms specify the amplitude floor *a*_min_, node response scale *c*_*i*_, reference input *L*_0_, and node load sensitivity *l*_*i*_. The phase response used a lagged gradient state,

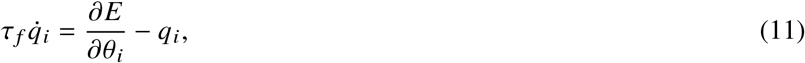

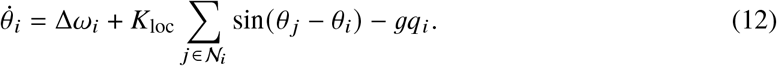

The state *q*_*i*_ is a low-pass-filtered phase gradient with response time *τ*_*f*_; Δ *ω*_*i*_ is the intrinsic frequency offset, *K*_loc_ is the nearest-neighbor coupling strength, *N*_*i*_ is the neighbor set, and *g* is the gradient-response gain. Common phase velocity was removed because Eq. (2) depends only on relative phase. The speed cap *v*_max_ limits the largest absolute relative velocity. The finite response time and speed cap therefore limit how quickly the phases can approach a moving target. Time was measured in nominal HSO cycles, with a primary integration step of 0.02 cycle. Numerical and structural checks varied the step, node count, boundary condition, input waveform, local attraction, initial phase, and speed limit.

### 2.4 Synthetic conditions and fixed comparators

The simulation program retained 98 discovery cases, 20 prespecified held-out cases, and a 45-case speed-cap challenge. The discovery cases were used to choose a response-reserve decision threshold; the held-out and challenge cases tested that threshold without refitting it. The held-out block contained four predefined families. Each family fixed one combination of node number (*N* = 6 or 9), between-node calcium-sensitivity variation (standard deviation 0.18 or 0.32), local attraction (0.015 or 0.04), population seed, and input waveform (triangular or asymmetric). Evaluating each family at environmental periods of 1.25, 2.5, 4, 6.5, and 10 cycles gave 20 cases. The speed-cap challenge crossed five environmental periods, three lags, and three speed limits.

The principal fixed comparator was the lowest evaluation-window result among 13 fixed candidates: an equal polygon and 12 configurations constructed from the initial amplitude vector. Each candidate remained constant and was constructed solely from equal-polygon geometry or the initial amplitude vector. Choosing the lowest outcome across the bank deliberately favored the fixed comparison: the dynamic response had to outperform the best fixed result found after evaluating all 13 candidates. Additional sensitivities used fixed phases optimized over the full evaluation trajectory for either mean *E* or CVaR_0.95_ (*E)*. Other controls used phase trajectories deliberately detached from the current amplitude state, including circular shifts, block permutations, time reversal, periodic motion, a mean-reverting stochastic trajectory with temporal correlation (Ornstein–Uhlenbeck motion), and a schedule transferred from training cases.

### 2.5 Tests of node-specific amplitude information

After the original response-surface analysis, I used the prespecified 20-case block to test whether the phase response requires node-specific amplitude assignment. Before calculating these input-comparison results, I fixed the four input drivers, gain grid, comparison rules, and directional criteria. The true amplitude matrix in Eq. (2) remained unchanged; only the amplitude information supplied to the response in Eqs. (11) and (12) varied. This controlled removal or reassignment of input information is often called an ablation test. The design kept the signal being evaluated identical while asking whether the mathematical phase-response rule needed to know which amplitude belonged to which node. The four drivers were the correctly aligned node-specific amplitudes, a node-independent root-mean-square amplitude, a one-node cyclic reassignment, and reversed node order. The two reassignments retained every amplitude sample and its temporal structure. Gain multipliers were 0, 0.25, 0.5, 1, 2, and 4.

The declared phase-motion cost was

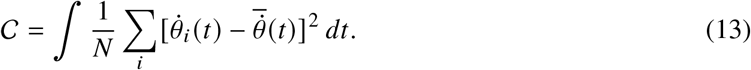

Thus, *C* accumulates the mean squared relative phase velocity and quantifies how rapidly and extensively the nodes rearrange their phases. Same-parameter cyclic and reversed drivers supplied the primary assignment controls because their motion cost matched the aligned driver numerically. The amplitude-blind driver served as a target-free control and commonly remained close to its equal-polygon initial state. Secondary comparisons selected the closest cost independently of outcome, or selected the lowest residual among candidates with *C*_control_ ≤ 1.05 *C*_aligned_.

The empirical assignment analysis retained the gain and lag selected within each leave-one-cell-out cross-validation fold[20]. In each fold, six cells selected the parameters and the remaining cell supplied an independent evaluation. For that held-out cell, I permuted only the five amplitude histories supplied to the phase response and evaluated *E* with the original node order. All 5! = 120 assignments were evaluated at the same selected parameters. Assignment rank was calculated within each cell, leaving seven independent biological units.

To measure the severity of an assignment error, I defined its mean serial displacement as

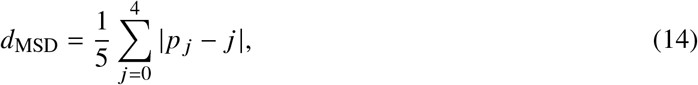

where *p*_*j*_ is the controller node receiving the amplitude history from observed node *j* . Within each cell, Spearman rank correlation related *d*_MSD_ to *Q*_0.95_ (*E* − *E*_min_); a positive value means that residuals tend to increase as assignments move farther from their observed positions. The seven cell-level correlations summarized the consistency and range of the graded assignment effect.

### 2.6 Response-reserve diagnostic

Response reserve was summarized by

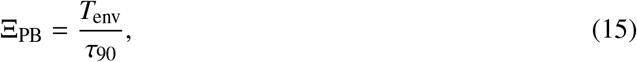

where *T*_env_ is the environmental period. To obtain *τ*_90_, I froze the amplitude vector at the time of maximum dynamic *E*, reset the lag state, and measured the time required for *E* to complete 90% of its decline toward the median over the final 20% of the relaxation trace. A large Ξ_PB_ means that the target changes over many response times and gives the phases ample time to adjust; a small value means that the target moves again before relaxation is complete. The ratio is a trajectory-derived diagnostic of the tested response; physiological interpretation requires independent calibration.

Dynamic advantage was

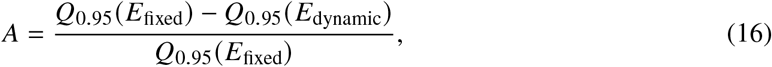

with *A >* 0 denoting a dynamic win. An increasing decision threshold was fitted in the 98 discovery cases by maximizing balanced accuracy and then applied unchanged to the 20 held-out and 45 challenge cases. Discrimination was assessed by the area under the receiver-operating-characteristic curve (AUC)[21] and balanced accuracy, the mean of the correct-classification rates for dynamic and fixed wins. An AUC of 0.5 indicates chance ranking and an AUC of 1 indicates perfect ranking of dynamic wins above fixed wins. A separate crossover analysis asked whether scaling by *τ*_90_ reduced between-family log-scale dispersion by at least 25%. Here, a crossover is the interpolated point where dynamic and fixed performance are equal, and log-scale dispersion measures how widely that point varies among condition families. Concentration of the normalized crossovers around one is termed collapse. Keeping these criteria separate allowed a useful classifier to coexist with family-dependent transition locations.

### 2.7 Signal processing and empirical replay

For the amplitude-versus-period analysis, each local trace was filtered around the cell-specific HSO peak ±1.0 Hz. Successive 2*π* phase wraps of the analytic signal defined cycle duration, and sample coefficients of variation (standard deviation divided by the mean) quantified amplitude and duration variability. Each cell was summarized by the median of its five trajectory-wise amplitude-CV/period-CV ratios. Thirty-six sensitivity settings crossed three neighboring 8.0-s HSO windows, two band widths, two cycle definitions, and three variability summaries.

The observable analysis first filtered each set of five traces in the 3.5–25-Hz HSO range. A third-order zero-phase Butterworth filter and Hilbert transform then supplied 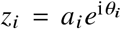 within the cell-specific HSO band. I compared *E* with a directly calculated, one-cycle-window power ratio of the summed real HSO-band traces. Within-cell correlation and absolute discrepancy quantified the agreement.

For replay, measured analytic amplitudes served as the model inputs. They were resampled every 0.02 cycle by past-value hold, which keeps the most recent available value until the next sample, smoothed by a one-sided exponential moving average with a primary time constant of 0.25 cycle, and normalized to unit row mean. Phases began from an equal polygon, and the relative-phase speed limit was 0.75 rad cycle^−1^. I used leave-one-cell-out selection: six cells selected gain from {10, 30, 60, 120} and lag from {0.04, 0.08, 0.16, 0.32} cycle by their median *Q*_0.95_ (*E* − *E*_min_), and the selected pair was then evaluated once on the seventh cell. A two-cycle initialization period preceded evaluation. Repeating this process held out each cell in turn and kept its outcome outside parameter selection. Fixed comparisons comprised the equal polygon, a configuration trained on the other six cells, and a personalized configuration based on the first two cycles of the held-out record. Circularly shifted amplitude histories preserved the same samples while breaking their timing relationship with the response. Smoothing and input-delay sensitivities tested time constants of 0.10 and 0.50 cycle and supplied-amplitude delays up to 0.50 cycle.

To reduce reliance on future samples, a separate sensitivity estimated amplitude using only the present and past signal. It replaced the zero-phase Hilbert envelope with a third-order forward-only band-pass filter followed by a one-sided exponential moving average of the squared signal and a square-root transform. The cell-specific dominant frequency and ±1-Hz band were retained from the primary analysis, and the controller parameters were kept at their original fold selections. This construction isolates the cost of one-sided amplitude estimation while preserving direct comparison with the primary replay.

The earlier multiband reconstruction used the same measured slow component and local HSO amplitudes in dynamic and fixed arms, while changing only relative HSO phase. Aggregate suppression in frequency band *b* was

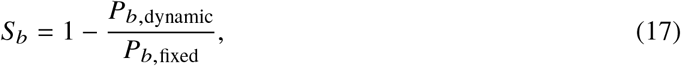

where *P*_*b*_ is the power of the summed band-limited length signal. Thus, *S*_*b*_ = 1 denotes complete suppression, *S*_*b*_ = 0 denotes unchanged power, and a negative value denotes an increase. Three slow bands and three HSO half-widths tested selectivity. A broad-band check compared unfiltered synthetic HSO input with 2.5–25-, 3.5–25-, and 4–25-Hz outputs and also summed the local oscillatory length-signal powers. The broad-band capture ratio was the captured broad-band power in the dynamic reconstruction divided by its fixed-reconstruction counterpart; a value of one means equal captured power. These robustness results are reported in Supplementary Figure S1.

### 2.8 Native phase compatibility

The native-phase analysis asked whether observed phase velocity followed the instantaneous direction that decreases *E*. Analytic amplitudes and unwrapped phases were processed with a Savitzky–Golay filter using a 0.25-cycle window and a third-degree local polynomial[22]; this local polynomial method smooths the signals and estimates phase velocity while preserving short-timescale waveform structure. Common phase velocity was removed. The phase contribution and directional alignment were

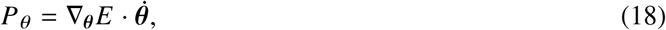

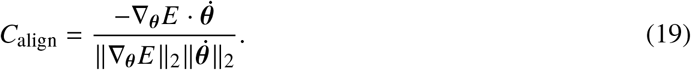

Negative *P*_*0*_ denotes instantaneous reduction of the length-signal residual. The normalized alignment *C*_align_ ranges from −1 to 1: 1 indicates motion exactly along the predicted improving direction, 0 indicates orthogonality, and −1 indicates the opposite direction. Twelve circular shifts, spaced from 1/13 to 12/13 of the complete phase-velocity record, retained the same observed phase movements but paired them with different times. This tested whether the timing of native phase motion agreed with the modeled beneficial direction better than expected from the same motion shifted in time. The primary criterion required observed mean *P* _*θ*_ below the shift median in at least six cells together with a one-sided cell-level Wilcoxon signed-rank *P <* 0.05[23]. Smoothing sensitivities used 0.10 and 0.50 cycle.

### 2.9 Statistics and numerical reproducibility

Recording-based inference used one summary per cell. Exact sign probabilities, Wilcoxon signed-rank tests, and cell-level bootstrap intervals obtained by resampling the seven cells with replacement were applied where defined in advance. Simulation conditions were described by counts, medians, ranges, and fixed comparison criteria; biological *P* values were reserved for cell-level inference. Receiver-operating curves and balanced accuracy evaluated the response-reserve classifier.

Analyses were implemented in Python with NumPy and SciPy[24, 25]. Numerical checks compared the analytic gradients with finite-difference approximations, confirmed that *E* decreases along the fixed-amplitude gradient direction, verified the polygon lower bound, and tested the time-varying balance in Eq. (6). After analysis, the full-precision condition- and cell-level values required for the principal numerical claims were retained. Supplementary Software provides a compact standard-library script that recalculates the 20-condition input-information comparison, the 163-condition response audit, the seven-cell replay summaries, the 840 within-cell node assignments, and the multiband sensitivity results; it also checks the central phasor equations.

## 3 Results

### 3.1 Amplitude heterogeneity creates a moving cancellation target

Figure 1A summarizes how the five measured sarcomere signals lead to the collective endpoint. Local HSO amplitude varied more strongly than fast-cycle duration in every cell. The cell-median amplitude- CV/period-CV ratio was 4.923, meaning that relative amplitude variability was about five times relative period variability. All seven paired cell summaries had the same direction (two-sided exact Wilcoxon signed-rank test, *P* = 0.015625; Figure 1B). The records contain the combination required for a moving cancellation target: appreciably changing vector lengths on a comparatively stable HSO timescale.

**Figure 1.**
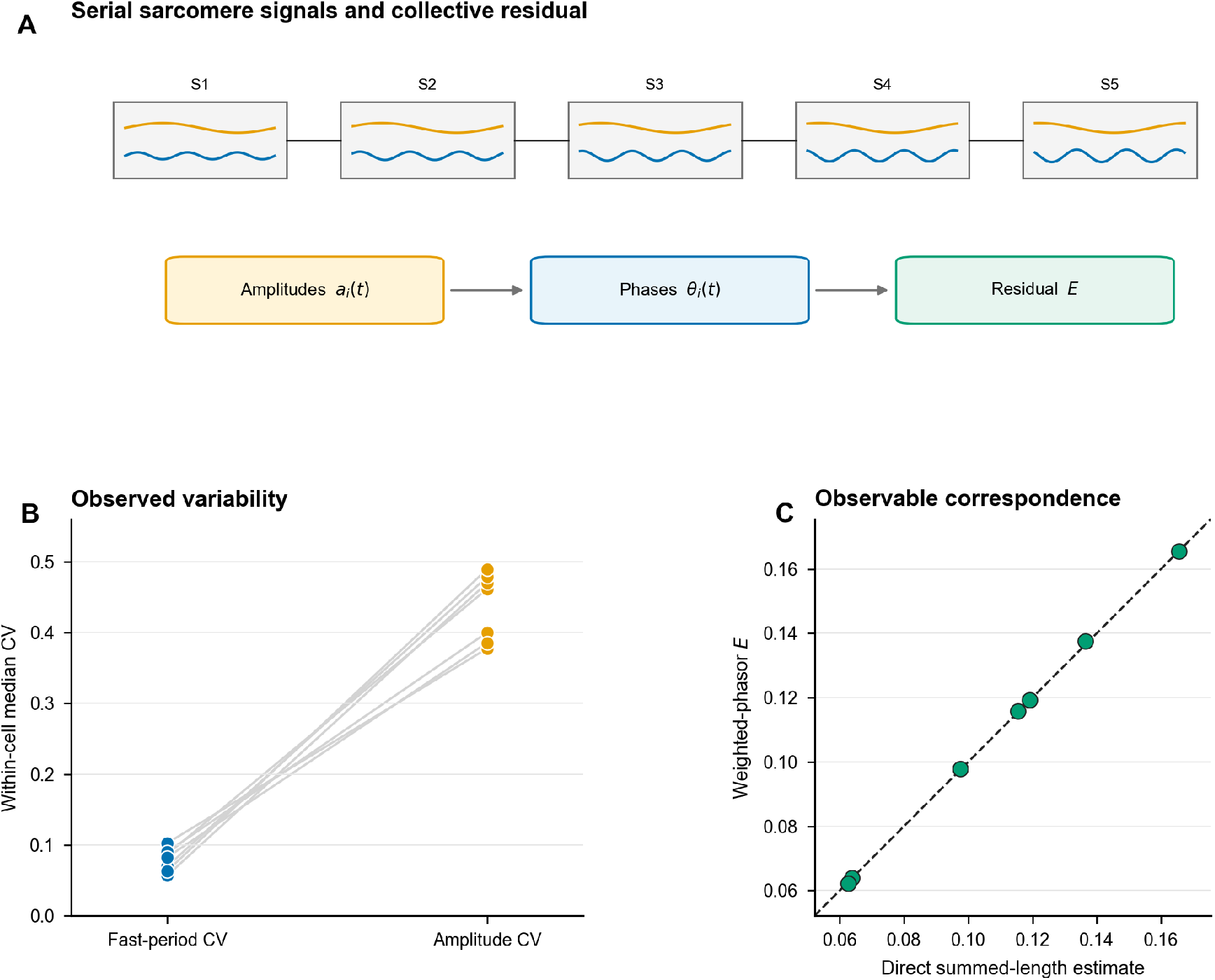
Empirical motivation and collective length-signal endpoint. (A) Schematic of five serial sarcomere signals. Orange and blue curves represent the slow and faster HSO components, respectively; the arrows summarize the reduced-model calculation from local signals to the collective endpoint. Relative HSO phase is the modeled variable, and *E* is the normalized residual of the summed HSO-band length signal. (B) HSO-amplitude variability exceeded fast-cycle variability in each of seven cells. Blue and orange circles denote period and amplitude variability, respectively, and gray lines pair the two summaries within a cell. The median trajectory-wise amplitude-CV/period-CV ratio was 4.923 (two-sided exact Wilcoxon signed-rank test, *P* = 0.015625). Amplitudes changed appreciably while the fast oscillation period remained comparatively stable. The cell is the independent unit. (C) The weighted-phasor residual followed the directly calculated one-cycle normalized power ratio of the summed real HSO-band signal. Each green circle represents one cell, and the dashed line marks equality. The cell-median within-cell correlation was 0.9971, and the cell-median 95th-percentile absolute discrepancy was 0.01371. This agreement supports use of *E* as the model endpoint for the recorded collective length signal. Panels B and C reanalyze the previously reported seven-cell series.

The weighted-phasor residual closely represented the directly summed HSO-band length signal. Across the seven cells, the median within-cell correlation between *E* and the one-cycle normalized summed-signal power ratio was 0.9971. The cell-median 95th-percentile absolute discrepancy was 0.01371 (Figure 1C). In other words, the compact vector calculation reproduced the collective signal obtained by directly summing the measured HSO-band traces closely enough to retain a physical interpretation as uncancelled chain-length fluctuation.

As the relative pattern of local amplitudes changes, the vector-length pattern changes and the phase directions that best cancel it can shift. In Eq. (6), *J* measures how strongly relative-amplitude change moves the residual, whereas *D* measures how strongly realized phase motion reduces it. Successful tracking requires phase motion to compensate for target motion before the target changes again (Figure 2A,B).

**Figure 2.**
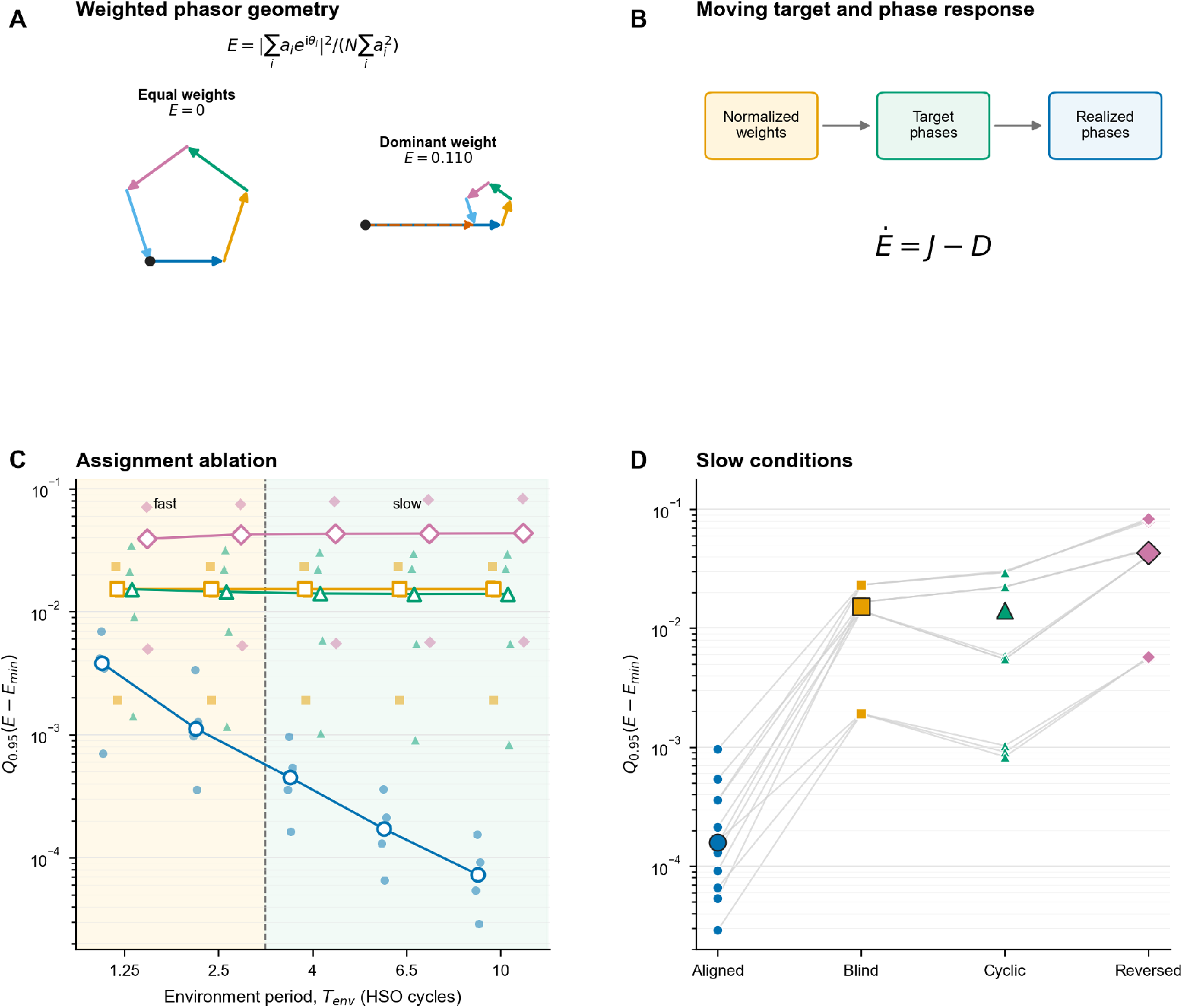
Correct node-amplitude alignment specifies the moving cancellation target. (A) Illustrative weighted-phasor geometry. The five colored arrows are the phasors 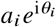: amplitude sets vector length and phase sets direction. A closed chain gives *E* = 0, whereas an open chain leaves a nonzero vector sum. The lower bound in Eq. (3) depends on whether the vectors can close a polygon. (B) Illustrative model sequence from normalized amplitudes to target phases and finite-lag realized phases. Changes in normalized amplitudes move the target phase configuration. Equation (6) separates the contribution of normalized-amplitude change, *J*, from the reduction generated by realized phase motion, . Response lag and the relative-phase speed limit restrict tracking. (C) Controlled input-information comparison across 20 prespecified conditions. Smaller values mean closer approach to the best cancellation allowed by the current amplitudes. Blue circles denote correctly paired input, orange squares amplitude-blind input, green triangles cyclic reassignment, and pink diamonds reversed order. Small filled symbols show individual conditions; large open symbols show period-specific medians, with colored lines connecting those medians. The vertical dashed line marks the fast–slow boundary. Pale orange and green backgrounds mark fast (*T*_env_ *<* 4 cycles) and slow (*T*_env_ ≥ 4 cycles) conditions, respectively. The correctly paired driver had lower upper-tail excess than each alternative in all 20 conditions. (D) Paired comparisons for the 12 conditions with *T*_env_ ≥ 4 cycles; gray lines connect the four input rules within the same model condition, symbol colors and shapes follow panel C, and large symbols show medians. Median *Q*_0.95_ (*E* − *E*_min_ ) was 0.0001585 for correctly paired input, 0.0152827 for amplitude-blind input, 0.0140523 for cyclic reassignment, and 0.0431218 for reversed order. Cyclic and reversed inputs preserved all amplitude samples and matched the declared motion cost numerically. Their larger residuals isolate the cost of incorrect node–amplitude pairing. The 20 model conditions are computational cases. of the current target.

### 3.2 Node-specific alignment determines which target is followed

Correctly paired amplitudes directed the model toward the lowest residual in all 20 input-ablation conditions. At the same gain, lag, speed limit, initial phase, and evaluation window, correctly paired input gave lower *Q*_0.95_ (*E* − *E*_min_) than the amplitude-blind, one-node cyclic, and reversed drivers in 20/20 conditions for each comparison (Figure 2C,D). In the 12 conditions with *T*_env_ ≥ 4 cycles, the respective medians were 0.0001585, 0.0152827, 0.0140523, and 0.0431218. Even the smallest control median was approximately 89 times the correctly paired median. The decisive information was the amplitude belonging to each particular node.

Node assignment explained the contrast with the permutation controls. Cyclic and reversed inputs preserved every amplitude sample and matched the correctly paired integrated squared relative-phase-speed cost to numerical precision (maximum absolute difference 4.996 × 10^−16^). The median paired-minus-cyclic difference in upper-tail excess was −0.0136516 over the 12 slow conditions. Matched samples and motion cost isolate correct node–amplitude pairing as the source of improvement.

### 3.3 Finite response capacity sets the range of dynamic advantage

The aligned response surpassed the strong fixed comparator when the amplitude target changed slowly. It won in 14/20 held-out conditions: 0/4 at an environmental period of 1.25 cycles, 2/4 at 2.5 cycles, and 4/4 at each of 4, 6.5, and 10 cycles (Figure 3A). This progression has a simple meaning: the fixed strategy led at the shortest target periods, whereas dynamic adjustment consistently led when four or more cycles were available. The median dynamic-minus-fixed *Q*_0.95_ (*E)* difference was −0.0004941 across all 20 conditions and −0.0015228 across the 12 slow conditions. Within-family interpolation placed the descriptive transition at 1.565–2.775 cycles.

**Figure 3.**
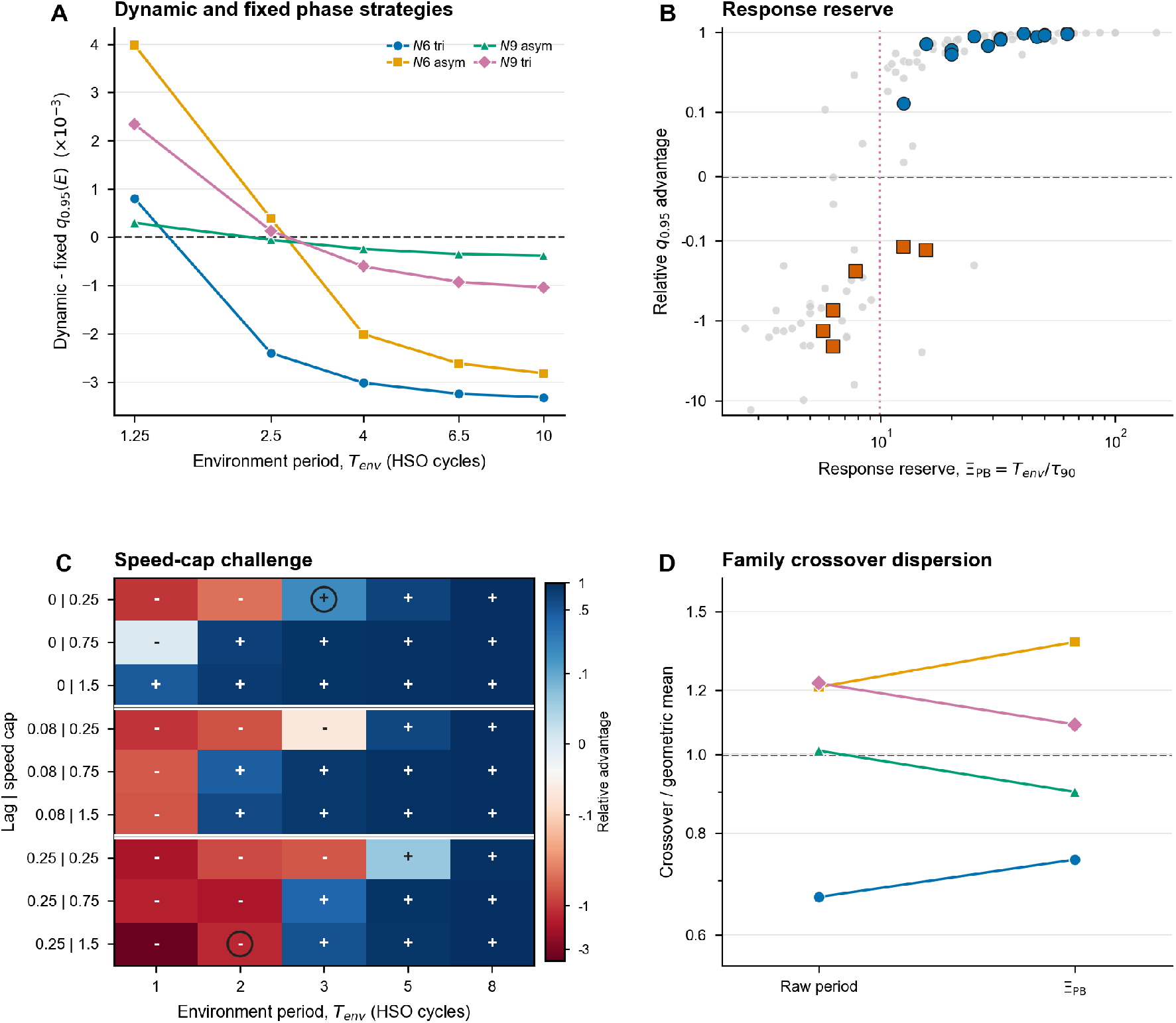
Finite response capacity defines the range of dynamic advantage. (A) Signed aligned-minus-fixed upper-tail residual for 20 held-out conditions. The fixed result is the lowest evaluation-window outcome among 13 candidates constructed solely from equal-polygon geometry or the initial amplitude vector. Negative values favor the dynamic response, and the horizontal dashed line marks equal performance. Colors and symbols identify four families defined by node number (N6 or N9) and triangular (tri) or asymmetric-pulse (asym) input. Dynamic phase adjustment won in 14/20 conditions: 0/4 at 1.25 cycles, 2/4 at 2.5 cycles, and 4/4 at 4, 6.5, and 10 cycles. (B) Response reserve, Ξ_PB_ = *T*_env_ /*τ*_90_, separated dynamic wins from fixed wins. Gray circles are the 98 discovery cases used to fit the threshold; blue circles and orange squares are held-out cases won by the dynamic and fixed strategies, respectively. The magenta dotted line is the discovery-fitted threshold, and the horizontal dashed line marks equal performance; positive relative advantage favors the dynamic strategy. Discovery AUC was 0.9709; the threshold 9.9026 gave held-out AUC 0.9762 and balanced accuracy 0.8333 (18/20). (C) Color encodes relative advantage (blue positive, red negative); ‘+’ and ‘−’ denote observed dynamic and fixed wins, respectively. Black rings mark the two cases misclassified by the unchanged threshold; row labels give lag (cycles) | speed cap (rad cycle^−1^), and horizontal separators divide the three lag blocks. The threshold gave AUC 0.9956 and balanced accuracy 0.9500 (43/45). (D) Colors and symbols identify the same four held-out families as in panel A. Each crossover is divided by the four-family geometric mean; lines connect the same family before and after normalization, and the dashed line at 1 marks the four-family geometric mean. Tighter clustering indicates stronger collapse. Between-family log-scale dispersion decreased by 6.81%, compared with the 25% criterion. Simulation conditions are computational cases. *τ*_90_ is calculated from a completed model trajectory and serves as a diagnostic of response capacity.

I next challenged the result with fixed phases optimized using the complete evaluation trajectory. Negative dynamic-minus-fixed values mean that dynamic adjustment left less residual. The two slow scenarios favored dynamic adjustment under both mean- and upper-tail-optimized fixed comparators, whereas the boundary and fast scenarios favored fixed phases. The signed differences ranged from −3.750 × 10^−4^ to +1.774 ×10^−3^. The fast conditions also had 2.65-fold higher median phase-motion cost and 14.6-fold higher aligned upper-tail residual than the slow conditions: the model was actively moving, but it was falling behind the target.

Response reserve asks how many response times fit within one period of target change. A larger value means more time to follow the target. In the discovery cases, Ξ_PB_ had AUC 0.9709, and the fitted threshold 9.9026 gave balanced accuracy 0.9466. Applied unchanged, the ratio correctly classified 18/20 held-out cases (AUC 0.9762; balanced accuracy 0.8333) and 43/45 speed-cap challenge cases (AUC 0.9956; balanced accuracy 0.9500), with 30 dynamic wins in the challenge (Figure 3B,C). These AUC values near 1 mean that response reserve almost perfectly ranked dynamic wins above fixed wins. Across all 163 conditions, the Spearman association between reserve and relative advantage was 0.9467.

The transition location retained a family dependence after normalization. Log-coordinate interpolation gave raw-period crossovers of 1.489–2.727 cycles and Ξ_PB_ crossovers of 9.303–17.234. Scaling reduced the between-family log-scale standard deviation from 0.2830 to 0.2638, a 6.81% reduction compared with the prespecified 25% criterion (Figure 3D). The ratio therefore provides strong ordering, while family-specific prediction also requires information about delay, required phase travel, and the geometry

### 3.4 Recorded amplitude histories support conditional model capacity

The recorded amplitude histories contained substantial variation on timescales accessible to the model. After one-sided smoothing, half of the spectral power in spatial amplitude-pattern variation lay at periods of about 11.0 HSO cycles or longer in the typical cell. Its autocorrelation half-decay was 1.808 cycles, the time over which pattern memory fell by half, and the median dwell of the largest-amplitude node was 1.617 cycles, the typical time before another node became largest. The median fraction of spectral power at periods at least as long as the conservative 2.775-cycle model crossover was 0.9851, with all seven cells above 0.50. Thus, most recorded amplitude variation lay on the slow side of the conservative model crossover, where the simulated response had time to follow.

Leave-one-cell-out replay gave a small residual above the instantaneous geometric bound in every held-out cell. All seven training folds selected gain 120 and lag 0.04 cycle, the edge of the defined grid. At these selections, median *Q*_0.95_ (*E* −*E*_min_) was 0.0008020673 for aligned replay, 0.077897998 for equal-polygon fixed phases, 0.062634190 for a generic fixed phase trained on the other six cells, and 0.100432570 for a personalized fixed phase constructed from the first two cycles of the held-out cell (Figure 4A). Because a smaller excess means closer approach to the best cancellation allowed by the measured amplitudes, the aligned median was about 1/78 of the lowest fixed-comparator median. Its advantage over all three fixed strategies in 7/7 cells was therefore both consistent and large on this normalized endpoint. It also remained lower than the median circular-shift control, 0.151353, in every cell. A 0.50-cycle supplied-amplitude delay increased the median excess to 0.019640 while retaining the direction relative to the equal fixed arrangement in every cell. Timely amplitude information was therefore part of the observed advantage.

**Figure 4.**
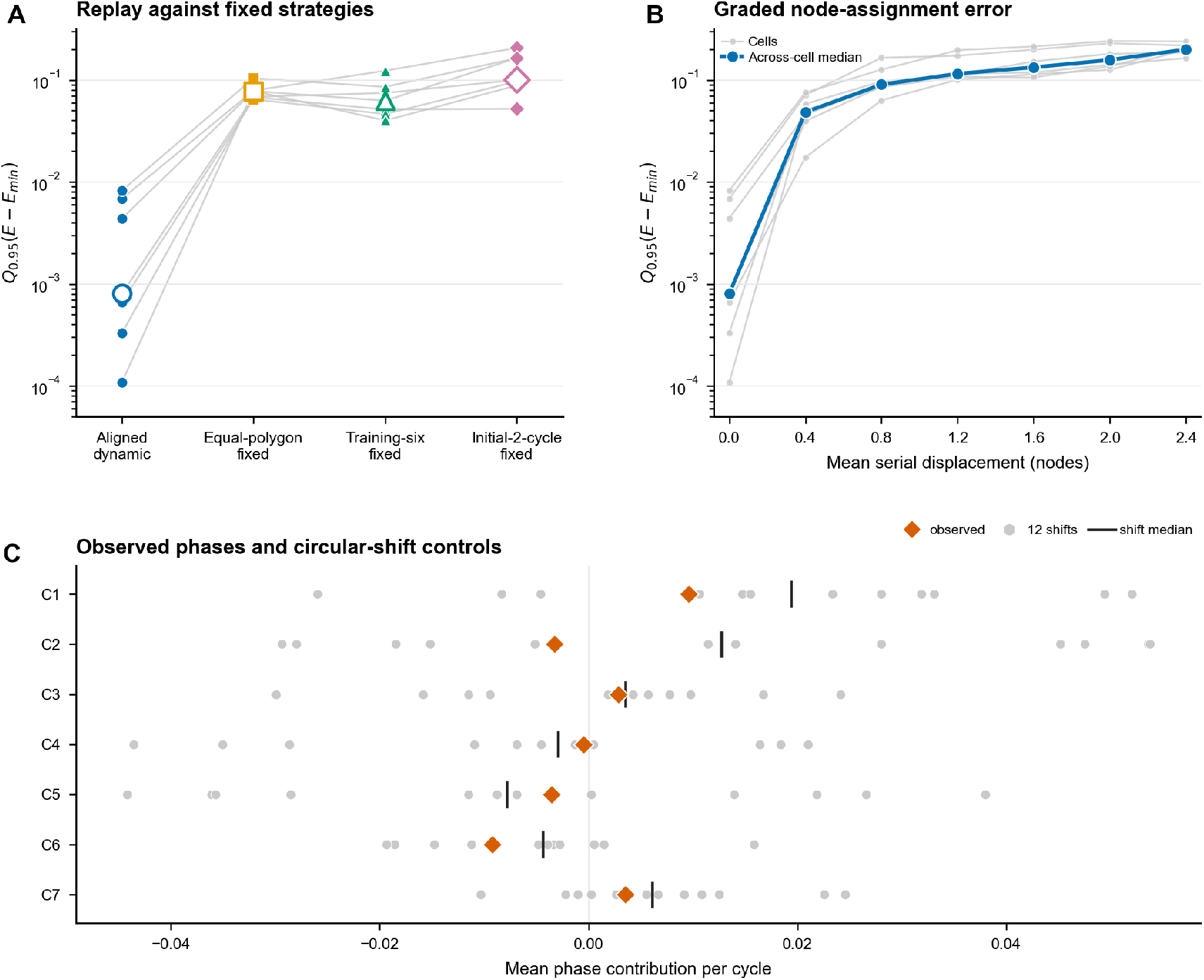
Sarcomere-derived replay supports assignment-dependent capacity. (A) Seven-fold leave-one-cell-out replay against three fixed strategies. Smaller values indicate closer approach to the instantaneous best-cancelling phase arrangement. Blue circles denote aligned dynamics, orange squares equal-polygon fixed phases, green triangles a generic fixed phase trained on the other six cells, and purple diamonds a fixed phase constructed from the first two cycles of the held-out cell. Small filled symbols are individual cells, gray lines pair the four strategies within each held-out cell, and large open symbols are seven-cell medians. Median *Q*_0.95_ (*E* −*E*_min_) was 0.000802, 0.07790, 0.06263, and 0.1004, respectively. Aligned replay was lower than each fixed strategy in all seven cells. (B) Graded node-assignment error at the unchanged fold-selected parameters. Mean serial displacement measures how far the five supplied amplitude histories move from their observed node positions. Thin gray lines and small gray circles show distance-bin medians for individual cells; the blue line and filled blue circles show the across-cell median. The within-cell Spearman correlation was positive in 7/7 cells, with a median of 0.568 and a range of 0.420–0.725. The identity mapping at zero displacement was the unique rank 1 among all 5! = 120 assignments in every cell. Larger spatial assignment errors caused progressively poorer cancellation. Assignments are within-cell controls, leaving *n* = 7 biological units. (C) Native-phase compatibility. Orange diamonds denote observed phase motion, gray points the 12 circular shifts, and black bars the shift medians. Negative values indicate phase motion that lowers *E*; the vertical line at zero marks no mean phase contribution. Observed phase motion was more residual-reducing than the shift median in 5/7 cells and more reducing than every shift in 0/7 cells; the one-sided cell-level Wilcoxon result was *P* = 0.1094. Amplitudes and phases were extracted offline.

A one-step extension of the selection grid confirmed the capacity result while showing that the original parameter search ended within an improving regime. The original gain 120 and lag 0.04 cycle reproduced the primary replay in 7/7 folds. Adding gain 240 and lag 0.02 cycle led every training fold to select the new boundary pair and lowered the held-out residual in 6/7 cells. The cell-median excess decreased by about 42%, from 0.000802 to 0.000465. Median relative phase-speed root-mean-square (RMS), a measure of typical phase-rearrangement rate, changed by about 0.05%, from 0.217277 to 0.217388 rad cycle^−1^. Median speed-limit occupancy, the fraction of evaluated samples at which the speed cap was active, increased by 0.25 percentage point, from 5.57% to 5.82% (Supplementary Figure S2A and Supplementary Table S1). The improvement therefore reflected a faster response with little change in total phase motion. I retained the original grid and leave-one-cell-out estimates as the primary analysis. The best setting still lay at the faster edge of the extended grid. These gain and lag values quantify tested computational capacity; physiological calibration belongs to the mechanochemical extension.

To reduce reliance on future samples, I repeated replay with an amplitude envelope calculated only from the present and past signal. At fixed response parameters, dynamic replay stayed below both the equal-polygon and training-six fixed comparators in 7/7 cells. The cell-median excess increased from 0.000802 with the offline envelope to 0.014379 with the forward-only envelope, compared with 0.082162 and 0.057631 for the two fixed comparators (Supplementary Figure S2B and Supplementary Table S2). The model advantage survived one-sided amplitude-envelope estimation, with lower tracking precision. This sensitivity evaluates the one-sided amplitude-estimation step while retaining the dominant frequency selected from the complete record. Online frequency selection remains for a fully real-time implementation.

Exhaustive reassignment showed that capacity depended on correct node-amplitude pairing. The observed identity mapping was the unique rank 1 among 120 assignments in each cell (Figure 4B). Across cells, median aligned excess was 0.000802, the median best nonidentity value was 0.02511, and the across-cell median of the seven within-cell nonidentity medians was 0.13484. All assignments contained the same five amplitude histories; only their node labels changed. Even the best wrong assignment was inferior, and a typical wrong assignment lost most of the cancellation advantage. A named cyclic reassignment had the same motion cost as the aligned replay to numerical precision and remained higher in all seven cells. The 120 assignments form within-cell controls; the biological sample remains seven cells.

Assignment errors also produced a graded loss. Within every cell, upper-tail excess increased with the mean serial displacement in Eq. (14). The Spearman correlation was positive in 7/7 cells, with a cell median of 0.568 and a range of 0.420–0.725 (Figure 4B). Performance worsened progressively as histories were assigned farther from their observed positions. The identity result is the optimum endpoint of this graded spatial effect.

The multiband reconstruction retained the same measured slow component and local HSO amplitudes in both arms. Median aggregate suppression was 0.9590 in the HSO band and −0.000626 in the slow band, with all seven cells meeting the defined selectivity criterion across all nine band definitions (Supplementary Figure S1). In plain terms, phase adjustment suppressed 95.90% of the summed fast-band power while leaving the slow signal essentially unchanged. The broad-band check gave median aggregate suppression of 0.9079 before output filtering and 0.9110 across 3.5–25 Hz. Dynamic-to-fixed ratios of summed local oscillatory length-signal power were 0.9999 and 0.9990, with a broad-band capture ratio of 0.9995. The amount of local oscillatory motion was therefore retained while its collective sum was reduced.

Native phase motion showed a small favorable average direction. At the primary smoothing scale, the cell-median phase contribution was −4.95 × 10^−4^ per cycle, so observed phase motion lowered *E* slightly on average, and median directional alignment was 0.0364 on the scale from −1 to 1. Observed motion reduced the residual more than the median circular shift in five cells and more than every shift in zero cells. The one-sided cell-level Wilcoxon result was *P* = 0.1094 (Figure 4C), and smoothing sensitivities retained this conclusion. This suggestive but subthreshold pattern identifies the predicted direction as a mechanochemical hypothesis for direct testing.

## 4 Discussion

The results identify two practical requirements for dynamic phase balancing. The spatial requirement is correct correspondence: the response must know which amplitude belongs to which node to identify the best-cancelling phase pattern. The temporal requirement is sufficient response capacity: phases must approach that pattern before changes in relative amplitudes move it elsewhere. Reassignment worsened cancellation even though the controls contained the same amplitude samples and produced the same total phase motion. Faster target motion then allowed a strong fixed pattern to perform better. Dynamic adjustment therefore adds value when both correct node-specific information and sufficient response time are available.

This formulation extends the earlier HSO studies in a specific direction. The 2020 mechanochemical study established amplitude modulation, relative period stability, and compensation of asynchronously changing sarcomere lengths and model tension[5]. The recent BPPB study characterized neighboring phase switching and the spatial reach of internal length redistribution in the same recordings[7]. Here, the weighted residual turns those observations into a moving-target problem. The assignment ablation identifies the local amplitude information that specifies the target, and the fixed comparison identifies the response range in which tracking adds value. The new contribution is an operating condition for a reduced model: aligned node amplitudes define the target, and finite response capacity determines its trackable range.

The exhaustive assignments give the spatial requirement a direct interpretation. The observed mapping remained best despite changing amplitudes, response lag, speed saturation, and finite evaluation windows. Performance then declined progressively as amplitude histories were assigned farther from their observed serial positions. The mathematics predicts an advantage for the correct mapping because the same node amplitudes define both the summed signal and its best-improvement direction. The informative result is that this advantage survived the specified finite response limits and comparison with every possible five-node assignment. A biological mechanism would need a physical signal, such as common tension or distributed strain, that conveys equivalent node-specific information through local mechanics.

The response-reserve analysis expresses the temporal requirement as a comparison of two clocks: how long the target takes to change and how long the phase response takes to relax. Their ratio separated dynamic and fixed wins accurately in held-out and challenge cases. The family-specific crossover positions identify delay, speed saturation, local attraction, initial geometry, and required phase travel as useful additional predictors. A future two-coordinate description could combine target drift with available phase travel. Prespecifying that definition before testing new conditions will protect the analysis from outcome-driven tuning.

Replay asked what the declared phase rule would do when driven by amplitude histories measured from real sarcomeres. The model residual closely followed the normalized power ratio calculated from the directly summed HSO-band signal, and much of the recorded amplitude variation occurred slowly enough to fall within the favorable simulated regime. Dynamic replay outperformed three fixed strategies in all seven cells, whereas delayed or forward-only amplitude input reduced its precision. Selection at the edge of the gain-lag grid shows that these values quantify the tested response capacity. Cellular gain and delay are targets for calibration in the next mechanochemical model, in which amplitude, phase, and mechanical load influence one another.

The native-phase comparison favored the modeled direction in 5/7 cells, with *P* = 0.1094 at the prespecified cell-level test. This pattern supports treating the present gradient as a beneficial direction predicted by the reduced model and the native response law as a mechanochemical hypothesis for direct testing. The earlier framework developed with Washio and colleagues offers a route to test its possible physical realization[5, 17, 26]. Small phase-indexed tension perturbations applied to identical copies of the same model state could measure the next-cycle phase shift and recovery time. Their relationship to perturbation phase would define a response kernel, which could then be compared with Eq. (4). This computational extension would use existing model states to connect the present operating range to actin– myosin cross-bridge and elastic mechanisms.

The study has clear boundaries. Biological inference rests on seven previously reported cells, whereas the larger simulation counts describe computational robustness. Every leave-one-cell-out fold selected the edge of the gain-lag grid, so the fitted values are operating coordinates for the tested model. The present model supplies node amplitudes to a global mathematical response and evaluates the summed HSO-band length signal. These limits define the scope of the claim: the present work establishes a length-signal capacity under measured amplitude input. A mechanochemical extension can calibrate cellular kinetics, derive a local physical response, and evaluate force, tension, mechanical work, ATP use, and period generation.

## 5 Conclusion

At each moment, node-specific amplitudes specify the phase arrangement that cancels the summed fast length changes most strongly. As their relative pattern changes, that arrangement can move. A phase response improves the collective signal when it receives the correct node-by-node information and moves quickly enough to follow the target; a strong fixed arrangement can perform better when the target changes too rapidly. Replay of sarcomere-derived amplitudes demonstrates this conditional capacity at the length-signal level. These findings turn asynchronous compensation into a testable mechanochemical question: can a serial sarcomere chain transmit the relevant amplitude information and adjust phase within the required time?

## Supporting information

Supplementary_Software

## 7 Author contributions

Seine A. Shintani designed the research, developed the model and analyses, analyzed the data, prepared the figures, and wrote the manuscript.

## 8 Data availability

The evidence data generated and/or analyzed during the current study are available from the corresponding author on reasonable request. Supplementary Software provides a compact standard-library reaggregation script and the full-precision condition- and cell-level fields needed to recalculate the reported summaries and check the core phasor equations.

## 9 Acknowledgements

This work was supported by JSPS KAKENHI Grant Number JP25K00269 (Grant-in-Aid for Scientific Research (C), project title: ‘‘Elucidation of Myosin Molecular Dynamics Associated with Sarcomere Morphological Changes in the Intracellular Environment’’).

During preparation of this work, the author used OpenAI Codex Desktop (version 26.818.5345.0; GPT-5, accessed August 2026) for language refinement and assistance with analysis and plotting code. The author reviewed and verified all code, numerical outputs, references, figures, and text and takes full responsibility for the publication.

