## Supplementary_Software for "Node-specific amplitude alignment and finite response capacity define dynamic phase balancing in a reduced sarcomere-chain model": README.txt

Focused Supplementary Software
==============================

Purpose
-------
This three-file package lets a reader recalculate the principal numerical
summaries reported in the manuscript and check the central phasor equations.
It does not make figures.

The package is deliberately limited to the evidence needed for the reported
claims.

Files
-----
README.txt
    This guide.

reproduce_results.py
    A single, standard-library Python script. It checks the core equations,
    recalculates the reported summaries, compares them with the full-precision
    reference values, prints PASS or FAIL, and returns a nonzero exit code if a
    check fails.

key_results.json
    Full-precision condition-level and cell-level values needed to identify
    the included conditions and recalculate the summaries, together with
    compact definitions and provenance.

Requirements and use
--------------------
Python 3.10 or later is sufficient. No third-party package is required.

From the extracted directory, run:

    python reproduce_results.py

To save the recalculated summary:

    python reproduce_results.py --output reproduced_summary.json

A successful run ends with PASS. The output file is optional and is not part
of the distributed archive.

What is recalculated
--------------------
The script performs the following checks without using saved summary values as
inputs to the calculations:

1. Core equations
   It evaluates normalized collective phasor energy, the polygon lower bound,
   the analytic phase gradient, and the sign of motion along the negative
   gradient.

2. Reduced-model information and response analyses
   It reaggregates 20 held-out information-ablation conditions and 163 response
   conditions (98 discovery, 20 held-out, and 45 speed-limit challenge cases).
   This includes dynamic-versus-fixed counts, the discovery-fitted response-
   reserve threshold, ROC AUC, balanced accuracy, Spearman correlation, and
   held-out crossover dispersion. The four full-trajectory fixed-comparator
   sensitivity cases are also checked.

3. Seven-cell replay
   It recalculates the cell medians and paired counts for amplitude and period
   variability, phasor-to-direct-power agreement, aligned replay, three fixed
   phase strategies, delayed input, the one-step response-boundary extension,
   forward-only amplitude processing, recorded timescales, multiband
   selectivity, broad-band power checks, and native-phase compatibility.
   Boundary improvements, band criteria, spectral checks, and native-phase
   comparisons are derived from their numerical values rather than from saved
   pass/fail flags. Exact signed-rank probabilities are calculated by
   enumeration for n=7.

4. Node assignment and band sensitivity
   It recalculates identity ranks, nonidentity summaries, and the within-cell
   relation between serial assignment distance and upper-tail residual for all
   7 x 120 node assignments. It also checks all 7 x 9 band-sensitivity rows.

Data dictionary
---------------
response_cases
    One row per computational response condition. T_env is the environmental
    period in nominal HSO cycles. tau90 is the frozen-input 90% relaxation time.
    The script calculates Xi_PB = T_env/tau90 and compares dynamic q95 with the
    best future-free fixed q95.

information_cases
    The 20 held-out model conditions. Each row contains the aligned,
    amplitude-blind, cyclic-node, reversed-node, and future-free fixed upper-
    tail values. Motion cost is integrated squared relative phase speed per
    node.

full_trajectory_cases
    Four sensitivity conditions comparing dynamic phases with fixed phases
    optimized over the complete amplitude trajectory.

empirical_cells
    One row per biological cell. Each value is derived from the previously
    reported recordings under the analysis definitions stated in the
    manuscript and Supplementary Information.

assignments
    The 120 permutations within each cell. An assignment such as 4-0-1-2-3
    states which measured amplitude history is supplied to each of the five
    model nodes. These permutations are controls within a cell, not biological
    replicates.

multiband_sensitivity
    Nine prespecified band definitions for each of the seven cells.

Scientific and data scope
-------------------------
The model-condition counts and node assignments are computational conditions.
The biological sample size is seven cells. The calculations test whether the
declared reduced phase rule can use correctly assigned amplitude histories;
they do not identify a natural phase controller or a mechanochemical cause.

The JSON begins from frozen, full-precision condition-level and cell-level
evidence. It does not contain the original experimental traces and does not
repeat signal extraction or every trajectory simulation. The recording source
used for the analyses has SHA-256

    1d1a272bc17d88ea28c6fd123959a18efe9176c6e97758246b6408e216c6d0b4

The evidence data are available from the corresponding author on reasonable
request, as stated in the manuscript. Figures, figure-generation code, and
internal development materials are outside the scope of this focused reader
package.
