## Supplementary_Software for "Node-specific amplitude alignment and finite response capacity define dynamic phase balancing in a reduced sarcomere-chain model": reproduce_results.py

```
#!/usr/bin/env python3
"""Recalculate the principal reported summaries without making figures."""
from __future__ import annotations

import argparse
import itertools
import json
import math
import statistics
import sys
from collections import defaultdict
from pathlib import Path
from typing import Iterable, Sequence


HERE = Path(__file__).resolve().parent
DEFAULT_DATA = HERE / "key_results.json"


def median(values: Iterable[float]) -> float:
    data = [float(value) for value in values]
    if not data:
        raise ValueError("median requires at least one value")
    return float(statistics.median(data))


def average_ranks(values: Sequence[float]) -> list[float]:
    order = sorted(range(len(values)), key=lambda index: values[index])
    ranks = [0.0] * len(values)
    start = 0
    while start < len(order):
        end = start + 1
        while end < len(order) and values[order[end]] == values[order[start]]:
            end += 1
        rank = 0.5 * (start + end - 1) + 1.0
        for position in range(start, end):
            ranks[order[position]] = rank
        start = end
    return ranks


def pearson(first: Sequence[float], second: Sequence[float]) -> float:
    if len(first) != len(second) or len(first) < 2:
        raise ValueError("correlation inputs must have equal length >= 2")
    first_mean = statistics.fmean(first)
    second_mean = statistics.fmean(second)
    numerator = sum(
        (x - first_mean) * (y - second_mean) for x, y in zip(first, second)
    )
    first_ss = sum((x - first_mean) ** 2 for x in first)
    second_ss = sum((y - second_mean) ** 2 for y in second)
    if first_ss <= 0.0 or second_ss <= 0.0:
        raise ValueError("correlation is undefined for a constant input")
    return numerator / math.sqrt(first_ss * second_ss)


def spearman(first: Sequence[float], second: Sequence[float]) -> float:
    return pearson(average_ranks(first), average_ranks(second))


def roc_auc(scores: Sequence[float], labels: Sequence[int]) -> float:
    positive = [score for score, label in zip(scores, labels) if label == 1]
    negative = [score for score, label in zip(scores, labels) if label == 0]
    if not positive or not negative:
        raise ValueError("ROC AUC requires both outcome classes")
    credit = 0.0
    for pos in positive:
        for neg in negative:
            credit += float(pos > neg) + 0.5 * float(pos == neg)
    return credit / (len(positive) * len(negative))


def balanced_accuracy(labels: Sequence[int], predictions: Sequence[int]) -> float:
    positives = [guess for truth, guess in zip(labels, predictions) if truth == 1]
    negatives = [guess for truth, guess in zip(labels, predictions) if truth == 0]
    if not positives or not negatives:
        raise ValueError("balanced accuracy requires both outcome classes")
    sensitivity = sum(guess == 1 for guess in positives) / len(positives)
    specificity = sum(guess == 0 for guess in negatives) / len(negatives)
    return 0.5 * (sensitivity + specificity)


def fit_increasing_threshold(
    scores: Sequence[float], labels: Sequence[int]
) -> tuple[float, float]:
    unique = sorted(set(float(value) for value in scores))
    if len(unique) < 2:
        raise ValueError("threshold fitting requires at least two score values")
    candidates = [math.nextafter(unique[0], -math.inf)]
    candidates.extend(
        0.5 * (left + right) for left, right in zip(unique[:-1], unique[1:])
    )
    candidates.append(math.nextafter(unique[-1], math.inf))
    best_threshold = candidates[0]
    best_score = -math.inf
    for threshold in candidates:
        predictions = [int(score >= threshold) for score in scores]
        score = balanced_accuracy(labels, predictions)
        if score > best_score + 1.0e-15:
            best_threshold = threshold
            best_score = score
    return best_threshold, best_score


def exact_signed_rank_p(differences: Sequence[float], alternative: str) -> float:
    nonzero = [float(value) for value in differences if value != 0.0]
    if not nonzero:
        return 1.0
    ranks = average_ranks([abs(value) for value in nonzero])
    observed = sum(rank for rank, value in zip(ranks, nonzero) if value > 0.0)
    possible = [
        sum(rank for rank, positive in zip(ranks, signs) if positive)
        for signs in itertools.product((False, True), repeat=len(ranks))
    ]
    tolerance = 1.0e-12
    lower = sum(value <= observed + tolerance for value in possible) / len(possible)
    upper = sum(value >= observed - tolerance for value in possible) / len(possible)
    if alternative == "less":
        return lower
    if alternative == "greater":
        return upper
    if alternative == "two-sided":
        return min(1.0, 2.0 * min(lower, upper))
    raise ValueError("alternative must be less, greater, or two-sided")


def normalized_energy(amplitudes: Sequence[float], phases: Sequence[float]) -> float:
    if len(amplitudes) != len(phases) or len(amplitudes) < 2:
        raise ValueError("amplitudes and phases must have equal length >= 2")
    resultant = sum(
        amplitude * complex(math.cos(phase), math.sin(phase))
        for amplitude, phase in zip(amplitudes, phases)
    )
    denominator = len(amplitudes) * sum(value * value for value in amplitudes)
    return abs(resultant) ** 2 / denominator


def polygon_lower_bound(amplitudes: Sequence[float]) -> float:
    total = sum(amplitudes)
    residual = max(2.0 * max(amplitudes) - total, 0.0)
    denominator = len(amplitudes) * sum(value * value for value in amplitudes)
    return residual * residual / denominator


def normalized_energy_gradient(
    amplitudes: Sequence[float], phases: Sequence[float]
) -> list[float]:
    units = [complex(math.cos(phase), math.sin(phase)) for phase in phases]
    resultant = sum(
        amplitude * unit for amplitude, unit in zip(amplitudes, units)
    )
    denominator = len(amplitudes) * sum(value * value for value in amplitudes)
    gradient = [
        2.0 * amplitude * (unit.conjugate() * resultant).imag / denominator
        for amplitude, unit in zip(amplitudes, units)
    ]
    mean_gradient = statistics.fmean(gradient)
    return [value - mean_gradient for value in gradient]


def core_equation_checks() -> dict[str, float]:
    equal = [1.0] * 5
    in_phase = normalized_energy(equal, [0.0] * 5)
    splay = normalized_energy(equal, [2.0 * math.pi * index / 5 for index in range(5)])
    lower_bound = polygon_lower_bound([5.0, 1.0, 1.0, 1.0, 1.0])
    amplitudes = [1.0, 0.7, 1.4, 0.9, 1.2]
    phases = [0.2, 1.1, -0.8, 2.2, -2.4]
    analytic = normalized_energy_gradient(amplitudes, phases)
    epsilon = 1.0e-6
    numeric: list[float] = []
    for index in range(5):
        plus = phases.copy()
        minus = phases.copy()
        plus[index] += epsilon
        minus[index] -= epsilon
        numeric.append(
            (normalized_energy(amplitudes, plus) - normalized_energy(amplitudes, minus))
            / (2.0 * epsilon)
        )
    gradient_error = max(abs(a - b) for a, b in zip(analytic, numeric))
    descent_rate = sum(value * (-3.0 * value) for value in analytic)
    return {
        "in_phase_energy": in_phase,
        "splay_energy": splay,
        "asymmetric_polygon_bound": lower_bound,
        "gradient_max_absolute_error": gradient_error,
        "negative_gradient_energy_rate": descent_rate,
    }


def response_record(row: dict[str, object]) -> dict[str, object]:
    period = float(row["environment_period"])
    tau90 = float(row["frozen_tau90"])
    dynamic = float(row["dynamic_q95"])
    fixed = float(row["best_future_free_fixed_q95"])
    return {
        **row,
        "xi90": period / tau90,
        "relative_advantage": (fixed - dynamic) / fixed,
        "dynamic_win": int(dynamic < fixed),
    }


def response_subset_metrics(
    rows: Sequence[dict[str, object]], threshold: float
) -> dict[str, float | int]:
    scores = [float(row["xi90"]) for row in rows]
    labels = [int(row["dynamic_win"]) for row in rows]
    predictions = [int(score >= threshold) for score in scores]
    return {
        "case_count": len(rows),
        "win_count": sum(labels),
        "auc": roc_auc(scores, labels),
        "balanced_accuracy": balanced_accuracy(labels, predictions),
        "correct_count": sum(truth == guess for truth, guess in zip(labels, predictions)),
        "spearman": spearman(
            scores, [float(row["relative_advantage"]) for row in rows]
        ),
    }


def held_out_crossovers(rows: Sequence[dict[str, object]]) -> list[dict[str, float | str]]:
    grouped: dict[str, list[dict[str, object]]] = defaultdict(list)
    for row in rows:
        grouped[str(row["family_id"])].append(row)
    output: list[dict[str, float | str]] = []
    for family, family_rows in sorted(grouped.items()):
        ordered = sorted(family_rows, key=lambda row: float(row["environment_period"]))
        crossing: tuple[int, float] | None = None
        for index in range(len(ordered) - 1):
            left = float(ordered[index]["relative_advantage"])
            right = float(ordered[index + 1]["relative_advantage"])
            if left == 0.0:
                crossing = (index, 0.0)
                break
            if left * right < 0.0:
                crossing = (index, -left / (right - left))
                break
        if crossing is None:
            continue
        index, fraction = crossing

        def interpolate(field: str) -> float:
            left = float(ordered[index][field])
            right = float(ordered[index + 1][field])
            return math.exp(math.log(left) + fraction * (math.log(right) - math.log(left)))

        output.append(
            {
                "family_id": family,
                "raw_period": interpolate("environment_period"),
                "xi90": interpolate("xi90"),
            }
        )
    return output


def assignment_distance(assignment: str) -> float:
    order = [int(value) for value in assignment.split("-")]
    if sorted(order) != list(range(len(order))):
        raise ValueError(f"invalid assignment: {assignment}")
    return sum(abs(index - value) for index, value in enumerate(order)) / len(order)


def multiband_selective(
    slow_suppression: float,
    hso_suppression: float,
    rules: dict[str, object],
) -> bool:
    return (
        hso_suppression
        >= float(rules["multiband_minimum_hso_suppression"])
        and abs(slow_suppression)
        <= float(rules["multiband_slow_preservation_tolerance"])
        and hso_suppression - slow_suppression
        >= float(rules["multiband_minimum_differential_suppression"])
    )


def spectral_escape_passes(
    row: dict[str, object], rules: dict[str, object]
) -> bool:
    unfiltered = float(row["unfiltered_aggregate_suppression"])
    broad = float(row["broad_aggregate_suppression"])
    local_low = float(rules["spectral_local_power_ratio_low"])
    local_high = float(rules["spectral_local_power_ratio_high"])
    capture_low = float(rules["spectral_capture_ratio_low"])
    capture_high = float(rules["spectral_capture_ratio_high"])
    return (
        unfiltered >= float(rules["spectral_minimum_aggregate_suppression"])
        and broad >= float(rules["spectral_minimum_aggregate_suppression"])
        and local_low <= float(row["unfiltered_local_power_ratio"]) <= local_high
        and local_low <= float(row["broad_local_power_ratio"]) <= local_high
        and capture_low <= float(row["broad_capture_ratio"]) <= capture_high
        and abs(broad - unfiltered)
        <= float(rules["spectral_maximum_filter_suppression_gap"])
    )


def calculate(payload: dict[str, object]) -> dict[str, object]:
    core = core_equation_checks()
    rules = dict(payload["analysis_rules"])

    information = list(payload["information_cases"])
    slow_minimum = float(rules["slow_environment_period_minimum_cycles"])
    slow = [
        row for row in information if float(row["environment_period"]) >= slow_minimum
    ]
    fast = [row for row in information if row not in slow]
    controls = ("unweighted_q95_excess", "cyclic_q95_excess", "reversed_q95_excess")
    information_summary = {
        "case_count": len(information),
        "slow_case_count": len(slow),
        "fast_case_count": len(fast),
        "aligned_better_than_each_control_count": sum(
            all(float(row["aligned_q95_excess"]) < float(row[field]) for field in controls)
            for row in information
        ),
        "dynamic_wins_total": sum(
            float(row["aligned_q95_excess"])
            < float(row["best_future_free_fixed_q95"])
            for row in information
        ),
        "dynamic_wins_slow": sum(
            float(row["aligned_q95_excess"])
            < float(row["best_future_free_fixed_q95"])
            for row in slow
        ),
        "dynamic_wins_fast": sum(
            float(row["aligned_q95_excess"])
            < float(row["best_future_free_fixed_q95"])
            for row in fast
        ),
        "slow_median_aligned_minus_unweighted": median(
            float(row["aligned_q95_excess"]) - float(row["unweighted_q95_excess"])
            for row in slow
        ),
        "slow_median_aligned_minus_cyclic": median(
            float(row["aligned_q95_excess"]) - float(row["cyclic_q95_excess"])
            for row in slow
        ),
        "cyclic_motion_max_absolute_difference": max(
            abs(float(row["aligned_motion_cost"]) - float(row["cyclic_motion_cost"]))
            for row in information
        ),
        "reversed_motion_max_absolute_difference": max(
            abs(float(row["aligned_motion_cost"]) - float(row["reversed_motion_cost"]))
            for row in information
        ),
    }

    response = [response_record(dict(row)) for row in payload["response_cases"]]
    discovery = [row for row in response if row["split"] == "discovery"]
    held_out = [row for row in response if row["split"] == "held_out"]
    challenge = [row for row in response if row["split"] == "speed_cap_challenge"]
    threshold, discovery_ba = fit_increasing_threshold(
        [float(row["xi90"]) for row in discovery],
        [int(row["dynamic_win"]) for row in discovery],
    )
    crossovers = held_out_crossovers(held_out)
    raw_crossovers = [float(row["raw_period"]) for row in crossovers]
    xi_crossovers = [float(row["xi90"]) for row in crossovers]
    raw_log_sd = statistics.stdev(math.log(value) for value in raw_crossovers)
    xi_log_sd = statistics.stdev(math.log(value) for value in xi_crossovers)
    response_summary = {
        "threshold": threshold,
        "discovery_balanced_accuracy": discovery_ba,
        "discovery": response_subset_metrics(discovery, threshold),
        "held_out": response_subset_metrics(held_out, threshold),
        "challenge": response_subset_metrics(challenge, threshold),
        "all": response_subset_metrics(response, threshold),
        "held_out_family_crossover_count": len(crossovers),
        "raw_period_crossover_min": min(raw_crossovers),
        "raw_period_crossover_max": max(raw_crossovers),
        "xi90_crossover_min": min(xi_crossovers),
        "xi90_crossover_max": max(xi_crossovers),
        "crossover_log_sd_reduction": 1.0 - xi_log_sd / raw_log_sd,
    }

    trajectories = {row["scenario"]: row for row in payload["full_trajectory_cases"]}
    trajectory_summary = {
        "case_count": len(trajectories),
        "success_and_representative_favor_dynamic": all(
            float(trajectories[name][field]) < 0.0
            for name in ("success", "development_representative")
            for field in (
                "dynamic_minus_mean_optimal_fixed",
                "dynamic_minus_cvar95_optimal_fixed",
            )
        ),
        "boundary_and_failure_favor_fixed": all(
            float(trajectories[name][field]) > 0.0
            for name in ("boundary", "failure")
            for field in (
                "dynamic_minus_mean_optimal_fixed",
                "dynamic_minus_cvar95_optimal_fixed",
            )
        ),
    }

    cells = list(payload["empirical_cells"])
    native_differences = [
        float(row["native_phase_contribution"])
        - float(row["native_shift_median_phase_contribution"])
        for row in cells
    ]
    empirical_summary = {
        "cell_count": len(cells),
        "amplitude_cv_median": median(float(row["amplitude_cv"]) for row in cells),
        "period_cv_median": median(float(row["period_cv"]) for row in cells),
        "amplitude_to_period_cv_ratio_median": median(
            float(row["amplitude_to_period_cv_ratio"]) for row in cells
        ),
        "amplitude_period_wilcoxon_two_sided_p": exact_signed_rank_p(
            [float(row["amplitude_cv"]) - float(row["period_cv"]) for row in cells],
            "two-sided",
        ),
        "power_ratio_pearson_r_median": median(
            float(row["power_ratio_pearson_r"]) for row in cells
        ),
        "power_ratio_q95_absolute_error_median": median(
            float(row["power_ratio_q95_absolute_error"]) for row in cells
        ),
        "aligned_q95_excess_median": median(
            float(row["aligned_q95_excess"]) for row in cells
        ),
        "equal_fixed_q95_excess_median": median(
            float(row["equal_fixed_q95_excess"]) for row in cells
        ),
        "training6_fixed_q95_excess_median": median(
            float(row["training6_fixed_q95_excess"]) for row in cells
        ),
        "personalized_fixed_q95_excess_median": median(
            float(row["personalized_fixed_q95_excess"]) for row in cells
        ),
        "shift_control_median_q95_excess_across_cells": median(
            float(row["shift_control_median_q95_excess"]) for row in cells
        ),
        "delay_0_5cycle_q95_excess_median": median(
            float(row["delay_0_5cycle_q95_excess"]) for row in cells
        ),
        "aligned_below_all_three_fixed_count": sum(
            all(
                float(row["aligned_q95_excess"]) < float(row[field])
                for field in (
                    "equal_fixed_q95_excess",
                    "training6_fixed_q95_excess",
                    "personalized_fixed_q95_excess",
                )
            )
            for row in cells
        ),
        "aligned_below_shift_median_count": sum(
            float(row["aligned_q95_excess"])
            < float(row["shift_control_median_q95_excess"])
            for row in cells
        ),
        "delay_below_equal_fixed_count": sum(
            float(row["delay_0_5cycle_q95_excess"])
            < float(row["equal_fixed_q95_excess"])
            for row in cells
        ),
        "primary_gain_values": sorted({float(row["primary_gain"]) for row in cells}),
        "primary_lag_cycle_values": sorted(
            {float(row["primary_lag_cycles"]) for row in cells}
        ),
        "boundary_primary_q95_excess_median": median(
            float(row["boundary_primary_q95_excess"]) for row in cells
        ),
        "boundary_extended_q95_excess_median": median(
            float(row["boundary_extended_q95_excess"]) for row in cells
        ),
        "boundary_primary_minus_extended_q95_excess_median": median(
            float(row["boundary_primary_q95_excess"])
            - float(row["boundary_extended_q95_excess"])
            for row in cells
        ),
        "boundary_extended_better_count": sum(
            float(row["boundary_extended_q95_excess"])
            < float(row["boundary_primary_q95_excess"])
            for row in cells
        ),
        "boundary_primary_phase_speed_rms_median": median(
            float(row["boundary_primary_phase_speed_rms"]) for row in cells
        ),
        "boundary_extended_phase_speed_rms_median": median(
            float(row["boundary_extended_phase_speed_rms"]) for row in cells
        ),
        "boundary_primary_speed_limit_occupancy_median": median(
            float(row["boundary_primary_speed_limit_occupancy"]) for row in cells
        ),
        "boundary_extended_speed_limit_occupancy_median": median(
            float(row["boundary_extended_speed_limit_occupancy"]) for row in cells
        ),
        "forward_dynamic_q95_excess_median": median(
            float(row["forward_dynamic_q95_excess"]) for row in cells
        ),
        "forward_equal_fixed_q95_excess_median": median(
            float(row["forward_equal_fixed_q95_excess"]) for row in cells
        ),
        "forward_training6_fixed_q95_excess_median": median(
            float(row["forward_training6_fixed_q95_excess"]) for row in cells
        ),
        "forward_below_both_fixed_count": sum(
            float(row["forward_dynamic_q95_excess"])
            < min(
                float(row["forward_equal_fixed_q95_excess"]),
                float(row["forward_training6_fixed_q95_excess"]),
            )
            for row in cells
        ),
        "spectral_median_period_cycles": median(
            float(row["spectral_median_period_cycles"]) for row in cells
        ),
        "acf_half_decay_cycles": median(
            float(row["acf_half_decay_cycles"]) for row in cells
        ),
        "top_node_dwell_median_cycles": median(
            float(row["top_node_dwell_median_cycles"]) for row in cells
        ),
        "fraction_period_ge_conservative_crossover_median": median(
            float(row["fraction_period_ge_conservative_crossover"])
            for row in cells
        ),
        "cells_fraction_period_ge_half": sum(
            float(row["fraction_period_ge_conservative_crossover"]) >= 0.5
            for row in cells
        ),
        "hso_band_suppression_median": median(
            float(row["hso_band_suppression"]) for row in cells
        ),
        "slow_band_suppression_median": median(
            float(row["slow_band_suppression"]) for row in cells
        ),
        "multiband_primary_gate_count": sum(
            multiband_selective(
                float(row["slow_band_suppression"]),
                float(row["hso_band_suppression"]),
                rules,
            )
            for row in cells
        ),
        "unfiltered_aggregate_suppression_median": median(
            float(row["unfiltered_aggregate_suppression"]) for row in cells
        ),
        "broad_aggregate_suppression_median": median(
            float(row["broad_aggregate_suppression"]) for row in cells
        ),
        "unfiltered_local_power_ratio_median": median(
            float(row["unfiltered_local_power_ratio"]) for row in cells
        ),
        "broad_local_power_ratio_median": median(
            float(row["broad_local_power_ratio"]) for row in cells
        ),
        "broad_capture_ratio_median": median(
            float(row["broad_capture_ratio"]) for row in cells
        ),
        "spectral_escape_gate_count": sum(
            spectral_escape_passes(dict(row), rules) for row in cells
        ),
        "native_phase_contribution_median": median(
            float(row["native_phase_contribution"]) for row in cells
        ),
        "native_better_than_shift_median_count": sum(
            float(row["native_phase_contribution"])
            < float(row["native_shift_median_phase_contribution"])
            for row in cells
        ),
        "native_better_than_all_shifts_count": sum(
            float(row["native_phase_contribution"])
            < float(row["native_shift_best_phase_contribution"])
            for row in cells
        ),
        "native_phase_wilcoxon_one_sided_p": exact_signed_rank_p(
            native_differences, "less"
        ),
    }

    assignment_rows = list(payload["assignments"])
    assignments_by_cell: dict[str, list[dict[str, object]]] = defaultdict(list)
    for row in assignment_rows:
        assignments_by_cell[str(row["cell"])].append(row)
    best_nonidentity: list[float] = []
    nonidentity_medians: list[float] = []
    distance_correlations: list[float] = []
    identity_unique_best = 0
    cyclic_motion_errors: list[float] = []
    for cell, rows in sorted(assignments_by_cell.items()):
        if len(rows) != 120:
            raise ValueError(f"{cell} has {len(rows)} assignments, expected 120")
        identity = next(row for row in rows if row["assignment"] == "0-1-2-3-4")
        nonidentity = [row for row in rows if row is not identity]
        identity_q95 = float(identity["q95_excess"])
        if sum(float(row["q95_excess"]) <= identity_q95 for row in rows) == 1:
            identity_unique_best += 1
        best_nonidentity.append(min(float(row["q95_excess"]) for row in nonidentity))
        nonidentity_medians.append(
            median(float(row["q95_excess"]) for row in nonidentity)
        )
        distance_correlations.append(
            spearman(
                [assignment_distance(str(row["assignment"])) for row in rows],
                [float(row["q95_excess"]) for row in rows],
            )
        )
        cyclic = next(row for row in rows if row["assignment"] == "4-0-1-2-3")
        cyclic_motion_errors.append(
            abs(float(identity["motion_cost"]) - float(cyclic["motion_cost"]))
        )
    assignment_summary = {
        "cell_count": len(assignments_by_cell),
        "assignment_count": len(assignment_rows),
        "identity_unique_rank1_cell_count": identity_unique_best,
        "best_nonidentity_median": median(best_nonidentity),
        "nonidentity_median_across_cells": median(nonidentity_medians),
        "distance_spearman_positive_cell_count": sum(
            value > 0.0 for value in distance_correlations
        ),
        "distance_spearman_median": median(distance_correlations),
        "distance_spearman_min": min(distance_correlations),
        "distance_spearman_max": max(distance_correlations),
        "cyclic_motion_max_absolute_difference": max(cyclic_motion_errors),
    }

    band_rows = list(payload["multiband_sensitivity"])
    by_setting: dict[tuple[float, float, float], list[dict[str, object]]] = defaultdict(list)
    for row in band_rows:
        key = (
            float(row["slow_low_hz"]),
            float(row["slow_high_hz"]),
            float(row["hso_halfwidth_hz"]),
        )
        by_setting[key].append(row)
    multiband_summary = {
        "row_count": len(band_rows),
        "setting_count": len(by_setting),
        "settings_passing_all_seven_cells": sum(
            len(rows) == 7
            and len({str(row["cell"]) for row in rows}) == 7
            and all(
                multiband_selective(
                    float(row["slow_suppression"]),
                    float(row["hso_suppression"]),
                    rules,
                )
                for row in rows
            )
            for rows in by_setting.values()
        ),
    }

    return {
        "core_equations": core,
        "information_ablation": information_summary,
        "response_reserve": response_summary,
        "full_trajectory_fixed": trajectory_summary,
        "empirical_replay": empirical_summary,
        "node_assignments": assignment_summary,
        "multiband_sensitivity": multiband_summary,
    }


def validate(summary: dict[str, object], expected: dict[str, object]) -> list[str]:
    tolerance = float(expected["absolute_tolerance"])
    errors: list[str] = []

    def exact(label: str, actual: object, target: object) -> None:
        if actual != target:
            errors.append(f"{label}: obtained {actual!r}, expected {target!r}")

    def close(label: str, actual: float, target: object) -> None:
        if not math.isclose(float(actual), float(target), rel_tol=0.0, abs_tol=tolerance):
            errors.append(f"{label}: obtained {actual:.16g}, expected {float(target):.16g}")

    core = summary["core_equations"]
    close("in-phase energy", core["in_phase_energy"], 1.0)
    if abs(float(core["splay_energy"])) > 1.0e-28:
        errors.append(f"splay energy is too large: {core['splay_energy']}")
    close("polygon lower bound", core["asymmetric_polygon_bound"], 1.0 / 145.0)
    if float(core["gradient_max_absolute_error"]) >= 1.0e-7:
        errors.append(f"gradient check failed: {core['gradient_max_absolute_error']}")
    if float(core["negative_gradient_energy_rate"]) >= 0.0:
        errors.append("negative-gradient direction did not reduce energy")

    info = summary["information_ablation"]
    exact("information case count", info["case_count"], expected["information_case_count"])
    exact(
        "aligned beats each information control",
        info["aligned_better_than_each_control_count"],
        expected["information_aligned_better_than_each_control_count"],
    )
    exact("held-out dynamic wins", info["dynamic_wins_total"], expected["dynamic_wins_held_out"])
    exact("slow dynamic wins", info["dynamic_wins_slow"], expected["dynamic_wins_slow"])
    exact("fast dynamic wins", info["dynamic_wins_fast"], expected["dynamic_wins_fast"])
    if float(info["cyclic_motion_max_absolute_difference"]) > 1.0e-12:
        errors.append("cyclic information control did not retain phase-motion cost")
    if float(info["reversed_motion_max_absolute_difference"]) > 1.0e-12:
        errors.append("reversed information control did not retain phase-motion cost")

    reserve = summary["response_reserve"]
    close("response threshold", reserve["threshold"], expected["response_discovery_threshold"])
    close("discovery AUC", reserve["discovery"]["auc"], expected["response_discovery_auc"])
    close("held-out AUC", reserve["held_out"]["auc"], expected["response_held_out_auc"])
    close(
        "held-out balanced accuracy",
        reserve["held_out"]["balanced_accuracy"],
        expected["response_held_out_balanced_accuracy"],
    )
    exact(
        "held-out correct count",
        reserve["held_out"]["correct_count"],
        expected["response_held_out_correct_count"],
    )
    close("challenge AUC", reserve["challenge"]["auc"], expected["response_challenge_auc"])
    close(
        "challenge balanced accuracy",
        reserve["challenge"]["balanced_accuracy"],
        expected["response_challenge_balanced_accuracy"],
    )
    exact(
        "challenge correct count",
        reserve["challenge"]["correct_count"],
        expected["response_challenge_correct_count"],
    )
    close("all-case Spearman", reserve["all"]["spearman"], expected["response_all_spearman"])
    close(
        "crossover log-SD reduction",
        reserve["crossover_log_sd_reduction"],
        expected["crossover_log_sd_reduction"],
    )

    trajectory = summary["full_trajectory_fixed"]
    exact("full-trajectory case count", trajectory["case_count"], 4)
    exact(
        "full-trajectory favorable signs",
        trajectory["success_and_representative_favor_dynamic"],
        True,
    )
    exact(
        "full-trajectory unfavorable signs",
        trajectory["boundary_and_failure_favor_fixed"],
        True,
    )

    empirical = summary["empirical_replay"]
    exact("biological cell count", empirical["cell_count"], 7)
    exact(
        "primary gain values",
        empirical["primary_gain_values"],
        [float(expected["primary_gain"])],
    )
    exact(
        "primary lag values",
        empirical["primary_lag_cycle_values"],
        [float(expected["primary_lag_cycles"])],
    )
    close(
        "amplitude-period Wilcoxon p",
        empirical["amplitude_period_wilcoxon_two_sided_p"],
        expected["amplitude_period_wilcoxon_two_sided_p"],
    )
    for key in (
        "aligned_q95_excess_median",
        "equal_fixed_q95_excess_median",
        "training6_fixed_q95_excess_median",
        "personalized_fixed_q95_excess_median",
        "delay_0_5cycle_q95_excess_median",
        "boundary_primary_q95_excess_median",
        "boundary_extended_q95_excess_median",
        "boundary_primary_minus_extended_q95_excess_median",
        "boundary_primary_phase_speed_rms_median",
        "boundary_extended_phase_speed_rms_median",
        "boundary_primary_speed_limit_occupancy_median",
        "boundary_extended_speed_limit_occupancy_median",
        "forward_dynamic_q95_excess_median",
        "forward_equal_fixed_q95_excess_median",
        "forward_training6_fixed_q95_excess_median",
        "hso_band_suppression_median",
        "slow_band_suppression_median",
        "unfiltered_aggregate_suppression_median",
        "broad_aggregate_suppression_median",
        "unfiltered_local_power_ratio_median",
        "broad_local_power_ratio_median",
        "broad_capture_ratio_median",
        "spectral_median_period_cycles",
        "acf_half_decay_cycles",
        "top_node_dwell_median_cycles",
        "fraction_period_ge_conservative_crossover_median",
        "native_phase_wilcoxon_one_sided_p",
    ):
        close(key, empirical[key], expected[key])
    exact("aligned below all fixed", empirical["aligned_below_all_three_fixed_count"], 7)
    exact("aligned below shift median", empirical["aligned_below_shift_median_count"], 7)
    exact("delayed below equal fixed", empirical["delay_below_equal_fixed_count"], 7)
    exact("boundary extension improved", empirical["boundary_extended_better_count"], 6)
    exact("forward below both fixed", empirical["forward_below_both_fixed_count"], 7)
    exact("timescale overlap cells", empirical["cells_fraction_period_ge_half"], 7)
    exact("primary multiband cells", empirical["multiband_primary_gate_count"], 7)
    exact("spectral escape cells", empirical["spectral_escape_gate_count"], 7)
    exact(
        "native below shift median",
        empirical["native_better_than_shift_median_count"],
        expected["native_better_than_shift_median_count"],
    )
    exact(
        "native below every shift",
        empirical["native_better_than_all_shifts_count"],
        expected["native_better_than_all_shifts_count"],
    )

    assignments = summary["node_assignments"]
    exact("assignment rows", assignments["assignment_count"], 840)
    exact("identity unique rank 1", assignments["identity_unique_rank1_cell_count"], 7)
    close(
        "best nonidentity median",
        assignments["best_nonidentity_median"],
        expected["assignment_best_nonidentity_median"],
    )
    close(
        "nonidentity median across cells",
        assignments["nonidentity_median_across_cells"],
        expected["assignment_nonidentity_median_across_cells"],
    )
    exact(
        "positive assignment-distance correlations",
        assignments["distance_spearman_positive_cell_count"],
        7,
    )
    close(
        "assignment-distance Spearman median",
        assignments["distance_spearman_median"],
        expected["assignment_distance_spearman_median"],
    )
    if float(assignments["cyclic_motion_max_absolute_difference"]) > 2.0e-14:
        errors.append("cyclic node assignment did not retain phase-motion cost")

    multiband = summary["multiband_sensitivity"]
    exact("multiband sensitivity rows", multiband["row_count"], 63)
    exact("multiband settings", multiband["setting_count"], 9)
    exact("multiband settings passing all cells", multiband["settings_passing_all_seven_cells"], 9)
    return errors


def print_summary(summary: dict[str, object], passed: bool) -> None:
    info = summary["information_ablation"]
    reserve = summary["response_reserve"]
    empirical = summary["empirical_replay"]
    assignments = summary["node_assignments"]
    multiband = summary["multiband_sensitivity"]
    print("Principal numerical checks")
    print(
        f"  Information controls: {info['aligned_better_than_each_control_count']}/"
        f"{info['case_count']} aligned cases lower than all three controls"
    )
    print(
        f"  Dynamic vs fixed: {info['dynamic_wins_total']}/20 overall; "
        f"{info['dynamic_wins_slow']}/12 slow; {info['dynamic_wins_fast']}/8 fast"
    )
    print(
        f"  Response reserve: threshold={reserve['threshold']:.7f}; "
        f"held-out AUC={reserve['held_out']['auc']:.4f}, "
        f"BA={reserve['held_out']['balanced_accuracy']:.4f}; "
        f"challenge AUC={reserve['challenge']['auc']:.4f}, "
        f"BA={reserve['challenge']['balanced_accuracy']:.4f}"
    )
    print(
        f"  Seven-cell replay medians: aligned={empirical['aligned_q95_excess_median']:.10f}; "
        f"equal fixed={empirical['equal_fixed_q95_excess_median']:.9f}; "
        f"training-six fixed={empirical['training6_fixed_q95_excess_median']:.9f}; "
        f"personalized fixed={empirical['personalized_fixed_q95_excess_median']:.9f}"
    )
    print(
        f"  Node assignment: identity rank 1 in "
        f"{assignments['identity_unique_rank1_cell_count']}/7 cells; "
        f"median distance rho={assignments['distance_spearman_median']:.3f}"
    )
    print(
        f"  Multiband sensitivity: {multiband['settings_passing_all_seven_cells']}/"
        f"{multiband['setting_count']} settings passed in all seven cells"
    )
    print("PASS" if passed else "FAIL")


def main() -> None:
    parser = argparse.ArgumentParser(
        description="Recalculate the principal reported numerical summaries."
    )
    parser.add_argument("--data", type=Path, default=DEFAULT_DATA)
    parser.add_argument("--output", type=Path)
    args = parser.parse_args()
    payload = json.loads(args.data.read_text(encoding="utf-8"))
    summary = calculate(payload)
    errors = validate(summary, dict(payload["expected_results"]))
    summary["validation"] = {"passed": not errors, "errors": errors}
    print_summary(summary, not errors)
    if errors:
        for error in errors:
            print(f"  - {error}", file=sys.stderr)
    if args.output is not None:
        args.output.write_text(
            json.dumps(summary, ensure_ascii=False, indent=2) + "\n",
            encoding="utf-8",
        )
    raise SystemExit(0 if not errors else 1)


if __name__ == "__main__":
    main()
```
