## Supplementary_Information for "Node-specific amplitude alignment and finite response capacity define dynamic phase balancing in a reduced sarcomere-chain model"

Seine A. Shintani

Department of Biomedical Sciences, College of Life and Health Sciences, Chubu University

Center for Mathematical Science and Artificial Intelligence, Chubu University

Institute for Advanced Research, Nagoya University

This document reports the band-specific reconstruction, the single-step leave-one-cell-out grid extension, the forward-only envelope sensitivity, and cell-level values for the fixed-phase and node-assignment comparisons. The biological sample is seven previously reported cardiomyocytes. Parameter settings and within-cell assignments are computational conditions or controls. Supplementary Software includes the full-precision fields required to recalculate the reported summaries and a compact standard-library reaggregation script.

### Supplementary Methods

##### Band-specific and broad-band reconstruction

The phase-only reconstruction used the measured 0.2–2.5-Hz slow component and the measured local HSO amplitudes in both the dynamic and fixed arms. Only the relative HSO phases differed. Aggregate suppression in band $b$ was

$$S_b = 1 - \frac{P_{b,\mathrm{dynamic}}}{P_{b,\mathrm{fixed}}}, \tag{1}$$

where $P_b$ is the power of the summed band-limited length signal. A value of 1 denotes complete suppression, 0 denotes unchanged power, and a negative value denotes an increase. Three definitions of the slow band and three HSO half-widths gave nine band combinations. The cell-level selectivity criterion required positive HSO-band suppression, an absolute slow-band effect below 0.05, and an HSO-minus-slow suppression difference above 0.50 in every band combination.

The broad-band check separated collective suppression from retention of the local oscillatory length signals. It compared dynamic and fixed aggregate signals before output filtering and after 3.5–25-Hz filtering, repeated the comparison for the sum of the five local signal powers, and calculated the broad-band capture ratio: dynamic captured power divided by the fixed counterpart. A ratio of one denotes equal captured power. Sensitivity bands with lower cutoffs of 2.5, 3.5, and 4.0 Hz shared an upper cutoff of 25 Hz. The measured amplitudes were identical in each paired reconstruction.

##### Strong fixed-phase comparators

The primary replay parameters were selected by leave-one-cell-out training and then applied once to the held-out cell. Three fixed strategies were evaluated over the same held-out window. The equal-polygon phases were fixed at equally spaced angles. The training-six configuration was optimized from 2,000 evenly spaced weight samples from each of the other six cells, using 24 seeded starts. The personalized fixed configuration was optimized from the first two cycles of the held-out cell; this preconditioning interval ended when the evaluation interval began. Each configuration remained fixed throughout evaluation. The endpoint was the 95th percentile of $E - E_{\min}$.

##### Single-step leave-one-cell-out grid extension

The primary selection grid contained gains $g \in \{10, 30, 60, 120\}$ and response lags $\tau_f \in \{0.04, 0.08, 0.16, 0.32\}$ HSO cycle. Every fold selected $g = 120$ and $\tau_f = 0.04$. A boundary audit added one value in each improving direction, $g = 240$

and $\tau_f = 0.02$, and repeated the same leave-one-cell-out selection. The smoothing time, speed limit, objective, training summaries, and held-out evaluation remained unchanged. The audit stopped after this single extension. Primary values were retained to preserve the prespecified primary analysis.

#### Forward-only envelope sensitivity

The primary empirical replay used a zero-phase Butterworth filter and Hilbert envelope to define the measured amplitude histories offline. The sensitivity analysis estimated amplitude from present and past samples using a third-order forward-only Butterworth band-pass filter, followed by a one-sided exponential moving average of the squared filtered trace and a square-root transform. Filter state was initialized from the first observed sample. The envelope time constant remained 0.25 HSO cycle, and values were resampled by past-value hold at 0.02-cycle intervals. The dominant frequency and its ±1-Hz band were retained from the complete-record estimate, and the original fold-selected phase-response parameters were reused. This test isolates the one-sided amplitude-estimation step. Real-time use also requires online frequency selection.

#### Assignment-distance summary

All 5! = 120 assignments of the five measured amplitude histories to the five model nodes were evaluated within each cell. Assignment distance was

$$d_{\mathrm{MSD}} = \frac{1}{5}\sum_{j=0}^{4} |p_j - j|, \tag{2}$$

where $p_j$ is the controller node receiving the amplitude history observed at node $j$. Spearman correlation related $d_{\mathrm{MSD}}$ to the assignment-specific 95th-percentile excess within each cell. These 120 assignments are within-cell controls; the seven cell-level correlations summarize consistency across the biological units.

### Supplementary Results

#### Collective suppression was concentrated in the HSO band

Median aggregate suppression was 0.9590 in the HSO band and −0.000626 in the slow band. All seven cells met the selectivity criterion in all nine band combinations (Supplementary Figure S1A). The broader check gave median aggregate suppression of 0.9079 before output filtering and 0.9110 after 3.5–25-Hz filtering. The corresponding dynamic-to-fixed ratios of summed local signal power were 0.9999 and 0.9990, and the median broad-band capture ratio was 0.9995 (Supplementary Figure S1B). The phase-only reconstruction therefore changed the summed fast length signal while retaining the component signals used to construct it.

#### The single-step grid extension improved six held-out cells

The primary candidate was reproduced in all seven folds. After one boundary step, every training fold selected the newly added pair, $g = 240$ and $\tau_f = 0.02$ cycle. Held-out upper-tail excess decreased in six cells and increased in cell 5. The cell median decreased from 0.000802 to 0.000465. Relative phase-speed RMS and speed-limit occupancy changed little at the cell median: 0.217277 to 0.217388 rad cycle$^{-1}$ and 5.57% to 5.82%, respectively (Supplementary Figure S2A and Supplementary Table S1). The result supports a faster-response capacity interpretation. The fitted settings quantify tested model capacity, and physiological calibration belongs to a mechanochemical model.

#### A forward-only envelope preserved the fixed-phase comparison

With the forward-only amplitude front end and unchanged controller settings, dynamic replay remained below the equal-polygon and training-six fixed values in all seven cells. Its median upper-tail excess increased from 0.000802 with the offline envelope to 0.014379, compared with 0.082162 for equal fixed phases and 0.057631 for the training-six fixed configuration under the forward-only front end (Supplementary Figure S2B and Supplementary Table S2). Thus, one-sided input processing reduced absolute tracking precision while retaining the comparative direction.

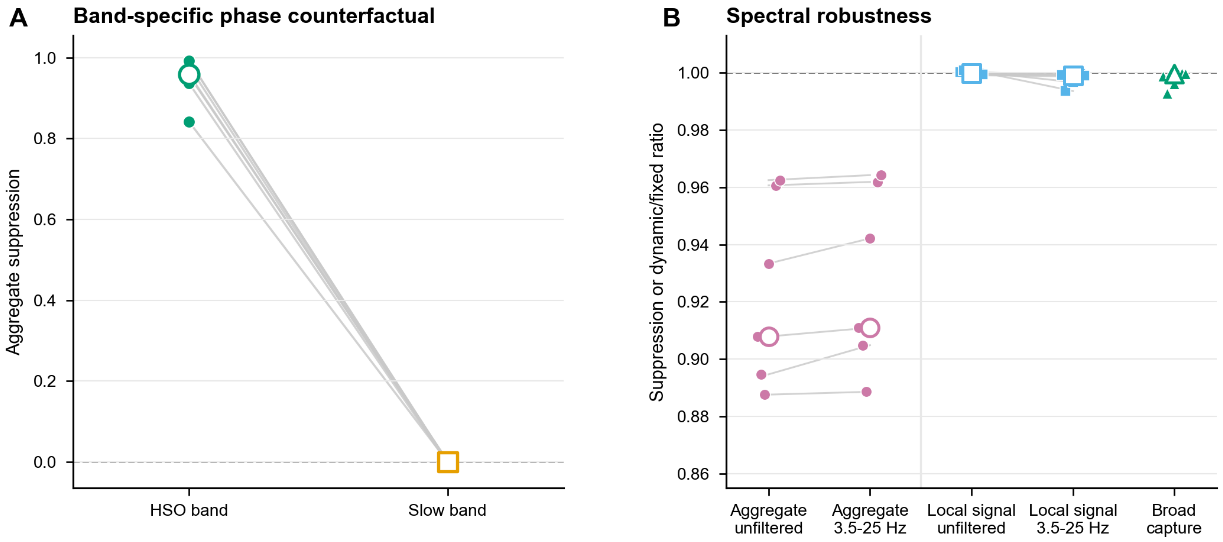


Figure S1: **Band-specific suppression and spectral robustness.** (A) Aggregate suppression for the HSO and slow bands in seven cells. Green circles and orange squares denote the HSO and slow bands, respectively. Zero denotes unchanged summed power and one denotes complete suppression. Small filled symbols are cell values, gray lines pair bands within each cell, and large open symbols are cell medians. The plotted points use the primary band definition; the all-nine statement summarizes the predefined band-width sensitivity. Median suppression was 0.9590 in the HSO band and −0.000626 in the slow band, and all seven cells satisfied the selectivity criterion under all nine definitions. (B) Broad-band checks in the same cells. Pink circles show aggregate suppression before filtering and after 3.5–25-Hz filtering; larger values mean a smaller summed signal. Blue squares show the dynamic-to-fixed ratio of summed local signal power, and green triangles show the ratio of broad-band captured power; for both ratios, one denotes unchanged local or captured power. Small filled symbols are cell values, gray lines pair unfiltered and filtered values from the same cell where both are shown, and large open symbols are cell medians. Median aggregate suppression was 0.9079 and 0.9110; median local signal-power ratios were 0.9999 and 0.9990; the median capture ratio was 0.9995.

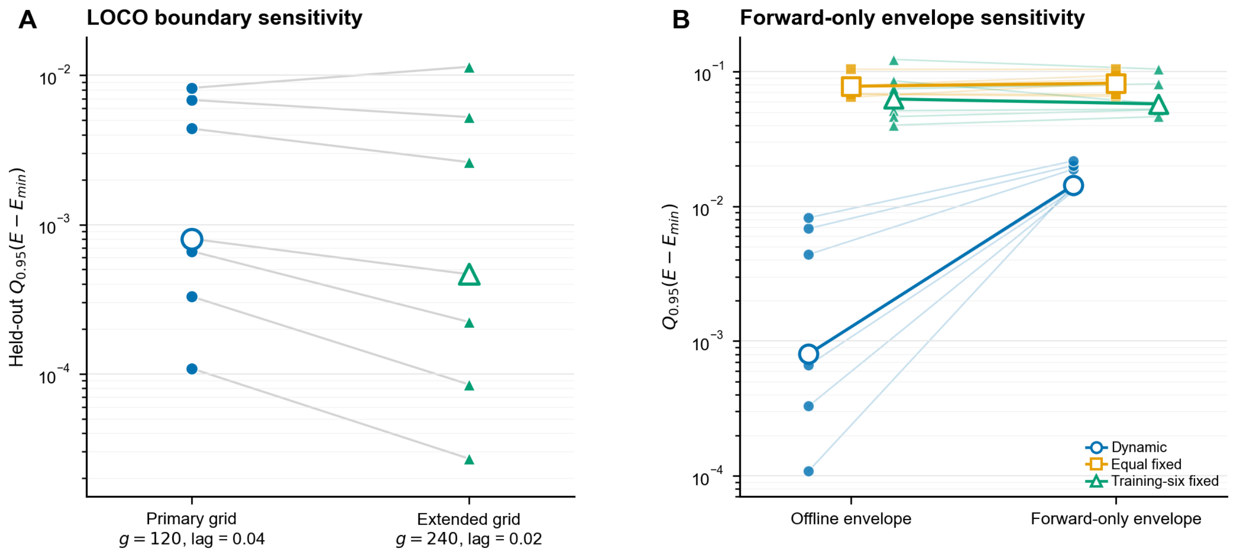


Figure S2: **Replay-boundary and forward-only-envelope sensitivities.** Smaller values in both panels mean closer approach to the instantaneous geometric bound. (A) Held-out upper-tail excess under the primary leave-one-cell-out grid (blue circles) and a single extension to $g = 240$ and lag = 0.02 cycle (green triangles). Gray lines pair the same cell, small filled symbols are cell values, and large open symbols are seven-cell medians. The extended setting lowered the endpoint in 6/7 cells; the median changed from 0.000802 to 0.000465. (B) Dynamic (blue circles), equal-fixed (orange squares), and training-six-fixed (green triangles) replay using the offline and forward-only amplitude envelopes. Thin lines pair the same cell within each strategy, small filled symbols are cell values, and large open symbols show medians. Under the forward-only envelope, dynamic replay remained lower than each fixed strategy in 7/7 cells, although its median upper-tail excess increased to 0.014379. This panel evaluates the one-sided amplitude-estimation step while retaining the dominant frequency estimated from the complete record.

### Supplementary Tables

#### Supplementary Table S1. Single-step leave-one-cell-out boundary audit

Primary settings were $g = 120$ and lag = 0.04 cycle; extended settings were $g = 240$ and lag = 0.02 cycle. $\Delta$ is primary minus extended held-out $Q_{0.95}(E - E_{\min})$, so positive values favor the extension. $\mathrm{RMS}_{\mathrm{pri}}$ and $\mathrm{RMS}_{\mathrm{ext}}$ are the root-mean-square relative phase speeds under the primary and extended settings, respectively, in rad cycle$^{-1}$. Occupancy is the percentage of evaluated samples at which the relative-phase speed limit was active.

| Cell | Primary | Extended | $\Delta$ | $\mathrm{RMS}_{\mathrm{pri}}$ | $\mathrm{RMS}_{\mathrm{ext}}$ | Occ. pri (%) | Occ. ext (%) |
|---|---|---|---|---|---|---|---|
| cell1 | 0.004394 | 0.002620 | 0.001774 | 0.2666 | 0.2682 | 18.75 | 19.25 |
| cell2 | 0.006846 | 0.005252 | 0.001594 | 0.2663 | 0.2658 | 13.15 | 12.15 |
| cell3 | 0.000802 | 0.000465 | 0.000337 | 0.1686 | 0.1698 | 5.19 | 5.59 |
| cell4 | 0.000329 | 0.000085 | 0.000244 | 0.2173 | 0.2174 | 1.57 | 0.81 |
| cell5 | 0.008234 | 0.011413 | −0.003179 | 0.2293 | 0.2458 | 13.00 | 15.78 |
| cell6 | 0.000663 | 0.000223 | 0.000440 | 0.1719 | 0.1731 | 5.57 | 5.82 |
| cell7 | 0.000108 | 0.000027 | 0.000081 | 0.1597 | 0.1592 | 1.12 | 1.12 |

#### Supplementary Table S2. Forward-only envelope sensitivity

Values are cell-level $Q_{0.95}(E - E_{\min})$. Offline dynamic uses the primary zero-phase/Hilbert amplitude envelope. The remaining columns use the forward-only filter and one-sided squared-signal envelope. RMS is the relative phase-speed RMS of the forward-only dynamic replay in rad cycle$^{-1}$.

| Cell | Dynamic offline | Dynamic forward | Equal fixed | Training-six fixed | RMS |
|---|---|---|---|---|---|
| cell1 | 0.004394 | 0.018911 | 0.078784 | 0.059383 | 0.487152 |
| cell2 | 0.006846 | 0.020147 | 0.093669 | 0.104497 | 0.474316 |
| cell3 | 0.000802 | 0.013861 | 0.065527 | 0.052685 | 0.491585 |
| cell4 | 0.000329 | 0.013019 | 0.067516 | 0.081120 | 0.477867 |
| cell5 | 0.008234 | 0.021756 | 0.103992 | 0.057631 | 0.482416 |
| cell6 | 0.000663 | 0.013241 | 0.082162 | 0.052120 | 0.474618 |
| cell7 | 0.000108 | 0.014379 | 0.087246 | 0.046344 | 0.482079 |

#### Supplementary Table S3. Strong fixed-phase comparisons

Values are cell-level $Q_{0.95}(E - E_{\min})$ under the primary fold-selected dynamic parameters. Dynamic replay was lower than each fixed strategy in every cell.

| Cell | Dynamic | Equal fixed | Training-six fixed | Initial-two-cycle fixed |
|---|---|---|---|---|
| cell1 | 0.004394 | 0.077898 | 0.062634 | 0.164359 |
| cell2 | 0.006846 | 0.078825 | 0.123271 | 0.206672 |
| cell3 | 0.000802 | 0.068055 | 0.051287 | 0.052232 |
| cell4 | 0.000329 | 0.068458 | 0.074621 | 0.100433 |
| cell5 | 0.008234 | 0.104122 | 0.085543 | 0.163983 |
| cell6 | 0.000663 | 0.064679 | 0.046345 | 0.096886 |
| cell7 | 0.000108 | 0.078120 | 0.040070 | 0.088351 |

#### Supplementary Table S4. Node-assignment distance summaries

The identity and all-assignment-median columns report cell-level $Q_{0.95}(E - E_{\min})$. Spearman $\rho$ relates mean serial displacement to assignment-specific upper-tail excess over all 120 within-cell assignments.

| Cell | Identity | All-assignment median | Spearman $\rho$ |
|---|---|---|---|
| cell1 | 0.004394 | 0.134735 | 0.657 |
| cell2 | 0.006846 | 0.199461 | 0.420 |
| cell3 | 0.000802 | 0.124202 | 0.563 |
| cell4 | 0.000329 | 0.149503 | 0.568 |
| cell5 | 0.008234 | 0.221271 | 0.464 |
| cell6 | 0.000663 | 0.113843 | 0.708 |
| cell7 | 0.000108 | 0.121906 | 0.725 |
